# An organoleptic survey of meads made with lactic acid-producing yeasts

**DOI:** 10.1101/445296

**Authors:** Carolyn Peepall, David G. Nickens, Joseph Vinciguerra, Matthew L. Bochman

## Abstract

We previously reported the isolation a suite of wild lactic acid-producing yeasts (LAYs) that enable “primary souring” during beer fermentation without the use of lactic acid bacteria. With sour meads gaining popularity in modern mead making, we were interested in exploring the same primary souring approach to traditional semi-sweet meads. In this study, we utilized 13 LAY strains to produce semi-sweet meads using a standardized batch of honey must to ensure consistent starting conditions. Thirteen 11-L batches of mead were prepared, and each was inoculated with one of the LAY strains, along with two control batches inoculated with champagne yeast. The initial pH and specific gravity were measured for each batch before inoculation. Traditional organic staggered nutrient addition was utilized for the first 72 h of fermentation with specific gravities being taken throughout the mead making process. Meads were racked, tasted, stabilized, cold crashed, bottled, and transported to the American Mead Maker’s Association 2018 Conference in Broomfield, Colorado. There, organoleptic surveys were conducted on these meads utilizing an array of tasters with varying levels of mead sensory analysis experience. The results of the sensory analysis, focusing on aroma and flavor, are discussed.

## 1. Introduction

Mead, and its origins, are the focus of many legends found deep within the lore of a variety of civilizations throughout the world. Mead is arguably the oldest alcoholic beverage known to humankind, preceding beer but perhaps not fruit-based wines, with mead evidence found in pottery dating to 6500-7000 BC in China (1) and in the Hindu texts of the Rigveda predating 1700 BC (2). The fact that both mead and fruit wines contain fewer ingredients and are easier to produce than beer, which requires the malting of grains to convert complex carbohydrates into readily fermentable sugars (3), also argues for the mead/wine before beer supposition.

The origin of mead is likely serendipitous as described by the Neolithic hypothesis (4): Honeycomb fell into a container of rain water (or rain water fell into a container of honey), wild yeasts initiated fermentation at mild ambient temperatures, and then a curious soul wandered by at the right time, tasted the elixir, and being ignorant of fermentation and alcohol, soon succumbed to the wiles of intoxication. No doubt a quest to replicate that magical elixir ensued, but it was not until the French chemist Louis Pasteur provided experimental evidence that yeast fermented sugars into ethanol in 1856 that alcoholic fermentation became more studied and understood (4-6). Until then, mead in its various forms became known for possessing sacred, mystical, and even healing properties. Receiving legendary notoriety as “ambrosia, or nectar, of the Gods” in ancient Greece, mead has notoriety as being the mythological source of the Norse god Odin’s strength when, as an infant, he drank mead from a goat’s udder (7).

Today, the fascination with mead and its mystical charm continues. Mead production is fast evolving with modern mead-making techniques riding on the coattails of craft brewing and distilling, rendering it easier to make and reducing fermentation times (8,9). More akin to wine making than beer brewing, mead has tapped into the creative minds of home and commercial mead makers alike (10). By replacing the grapes and grape juice typically found in wines with various honey varietals and adding yeast, water, and nutrients, traditional meads are born. Meads can vary from being light-bodied and low in alcohol to being sweeter, full bodied, and containing a higher alcohol by volume (ABV). They also vary in sweetness from being dry, semi-sweet, or sweet, which is not dissimilar from typical wine characteristics. Yeast choice has certain organoleptic influences on mead, impacting mouthfeel, aromas, and flavors (11). A plethora of adjunct additions (*e.g.*, fruits and spices) and wood aging techniques may also be used to make meads more complex. However, once again like wine, achieving the delicate balance between sweetness, alcohol, tannins, and acidity is the key to making an exceptional mead.

The Beer Judge Certification Program (BJCP) guide on mead styles (12) provides insight into how much creative license craft mead making provides. The Experimental Mead category provides the creative mead maker with almost endless options for barrel aging, fortification, and experimenting with microbes. It was with this creative inspiration that we sought to use heterofermentative lactic acid-producing yeast (LAY) to make meads and perhaps develop a novel Sour Mead category in the BJCP Mead Styles. As documented in the 2017 American Mead Makers Association (AMMA) Mead Industry Report, sour meads currently account for 13.2% of the mead styles being commercially produced (9). Current souring techniques for both beer and mead require either rapid kettle souring or slow techniques using mixed culture (lactic acid bacteria and yeast) fermentation (13), both presenting risk of infection across batches. Recently, the use of LAY has been suggested as an alternative to these techniques (14), being as rapid as kettle souring but reducing the risk for contamination of other batches because LAY are susceptible to traditional sanitation procedures. Here, we used a collection of 13 LAY strains to produce sour meads and assessed their perceived organoleptic properties by surveying a panel of mead sensory analysts at the 2018 AMMA Conference. The detailed fermentation and sensory results are presented below, and we conclude that LAY strains expand the options available for mead makers in producing sour meads.

## 2. Methods and materials

### 2.1. Strains, media, and culture other reagents

*S. cerevisiae* strain WLP715 (champagne yeast (15)) was purchased from White Labs (San Diego, CA). The LAY strains were provided by Wild Pitch Yeast (Bloomington, IN) and are described in (14). All yeast strains were routinely grown on yeast extract, peptone, and dextrose (YPD; 1% (w/v) yeast extract, 2% (w/v) peptone, and 2% (w/v) glucose) plates containing 2% (w/v) agar at 30°C and in YPD liquid culture at 30°C with aeration unless otherwise noted. All strains were stored as 15% (v/v) glycerol stocks at −80°C. Media components were from Fisher Scientific (Pittsburgh, PA, USA) and DOT Scientific (Burnton, MI, USA). All other reagents were of the highest grade commercially available.

### 2.2. Mead fermentation

Fifteen 11-L batches of traditional mead must, with an original gravity (OG) of 1.112 and initial pH of 4.13, were prepared using true source clover honey (Dutch Gold, Lancaster, PA) and spring water. Each was inoculated with approximately 1.85 × 10^10^ cell of one of the 13 LAY strain or the WLP715 control and degassed using a degassing wand attached to a standard electric drill. To assist with clarification, 22 g (1-3g/L) of dry Albumex Perl Bentonite (Max F. Keller GmbH, Mannaheim, Germany) was added to each batch 12 h before inoculation. Traditional organic staggered nutrient addition (TOSNA) (16) using a FermaidO (Lallemand Oenology, Canada) regimen and twice daily degassing were utilized to provide the yeast strains with sufficient and consistent nutrients, as well as oxygenation during the first 72 h of fermentation. The ambient and mead fermentation temperatures ranged from 17-20°C.

The meads were racked off the yeast 3 weeks after inoculation, and the specific gravity (SG) for each batch was measured using a hydrometer. At 6 weeks after inoculation, each batch was racked for a second time, and the SG was again recorded. Tastings were performed intermittently throughout the process. In the ninth week, a 6-g (2 g/3.7-L) potassium sorbate (LD Carlson Co., Kent, OH) 0.5-g (lg/7.4-L) potassium metabisulfite (LD Carlson Co., Kent, OH) combination was added to stabilize the meads. Fermfast DualFine Super Kleer (LD Carlson Co., Kent, OH) was added as a fining agent per the manufacturer’s instructions. Briefly, DualFine solution A1 (silica) was gently stirred into the mead and allowed to incubate for 1 h. Then, Dualfine solution A2 (chitosan) was gently stirred in for 30 s. The meads were cold-crashed by being incubated at an ambient temperature of −2.2 to 3°C for 36 h before bottling to aid in clearing. All mead making equipment was thoroughly cleaned and sanitized with Star San (Five Star Chemicals and Supply, Inc., Commerce City, CO.) prior to each step of the mead making process. The bottles were transported under refrigeration to the 2018 AMMA Conference in Broomfield, CO, to conduct tasting and sensory evaluations (below).

### 2.3. Sensory analysis

The goal of this experiment was to analyze the mead sensory characteristics produced by the 13 LAY strains utilizing a wide variety of taster sensory analysis experience levels, as well as variation in palate sophistication. The benefit of this, rather than only using expert sensory analysts, was to gather feedback from a population that would mimic the full range of craft mead consumers, from individuals tasting their first mead to long-term enthusiasts. The tasting occurred at the 2018 AMMA Mead Conference and Trade Show during the “AMMA Grant: Sour Mead Tasting and Feedback Session (https://mead-makers.org/2018-amma-mead-conference-and-trade-show/keynote-speaker-dr-patrickmcgovern/wednesday-march-14/#Sour). After a brief verbal introduction to the LAY sour mead project, sensory analysis feedback sheets (see Supplemental Materials for examples) were provided to tasters for all 13 batches of experimental mead. Tasters were asked to provide their feedback the flavor, aroma, and mouthfeel, of each mead. In this informal setting, the sensory volunteers were given as much time as possible to smell and taste the meads, as well as individually document their feedback on the sensory analysis sheets. A total of 490 sensory analysis feedback sheets were completed and gathered during the mass tasting session held at the 2018 AMMA Conference. However, it should be noted that not all tasters completed their surveys. For instance, seven out of 40 tasters of the mead fermented with strain YH25 failed to record their expertise level (see Supplemental Materials). We chose to include data from incomplete survey sheets because participants were not explicitly instructed to fill out every section of the surveys. Further, our goal was not to categorize our participants and the data from individual groups but simply to receive feedback from a broad range of tasters, from novices to experts.

### 2.4. Overall impression scoring

Overall, the meads were ranked on a scale from 1-5, with 5 being most favorable. The sums of these overall impression scores were then calculated, taking into account the score of each volunteer who tasted the mead. For instance, the mead fermented by YH77 received no scores of 5, three scores of 4, 23 scores of 3, 12 scores of 2, and two scores of 1. Multiplying the number of scores by the score value and summing the total yields an overall impression score of 0 + 12 + 69 + 24 + 2 = 107 for the YH77 mead. The average of these scores across all of the experimental meads was calculated to determine how individual meads ranked relative to the entire group. If 40 surveys were returned for a mead, each with the highest rank of 5, then the perfect mead would have received an overall impression score of 200.

### 2.5. Statistical analysis

All sensory analysis data from the feedback sheets were entered into Microsoft Excel spreadsheets. Statistics were determined using Microsoft Excel and/or GraphPad Prism.

## 3. Results

### 3.1. Fermentation performance

To compare the fermentative abilities of all 13 LAY strains to the WLP715 control, 170 L of honey must was prepared using clover honey and spring water, and 11 L was added to each of 15 individual fermenters. These vessels were then each inoculated with a different LAY or the control strain (x2) and allowed to ferment at ambient temperature (17-20°C). Samples of each batch were collected several times during fermentation to measure SG and collect tasting notes (Supplemental Table S1). Table 1 shows the specific gravity, attenuation, ABV, and pH information for the 15 meads, including the two control batches (C1 and C2) at the end of fermentation.

**Table 1.** Fermentation parameters.

| Strain | Species | OG<sup>a</sup> | FG | Attenuation (%) | ABV (%) | Initial pH | Terminal pH |
| --- | --- | --- | --- | --- | --- | --- | --- |
| WYP39 | *L. fermentati* | 1.112 | 1.068 | 37 | 5.8 | 4.12 | 3.24 |
| YH25 | *L. fermentati* | 1.112 | 1.062 | 42 | 6.56 | 4.12 | 3.33 |
| YH26 | *L. thermotolerans* | 1.112 | 1.070 | 35 | 5.6 | 4.12 | 3.13 |
| YH27 | *L. thermotolerans* | 1.112 | 1.034 | 68 | 10.2 | 4.12 | 3.15 |
| YH72 | *H. vineae* | 1.112 | 1.080 | 27 | 4.2 | 4.12 | 3.07 |
| YH73 | *L. thermotolerans* | 1.112 | 1.072 | 34 | 5.3 | 4.12 | 3.12 |
| YH77 | *L. fermentati* | 1.112 | 1.066 | 39 | 6.0 | 4.12 | 3.35 |
| YH79 | *L. thermotolerans* | 1.112 | 1.044 | 58 | 8.9 | 4.12 | 3.00 |
| YH81 | *L. thermotolerans* | 1.112 | 1.038 | 64 | 9.7 | 4.12 | 3.02 |
| YH82 | *W. anomalus* | 1.112 | 1.070 | 35 | 5.6 | 4.12 | 3.1 |
| YH109 | *L. thermotolerans* | 1.112 | 1.056 | 48 | 7.4 | 4.12 | 3.01 |
| YH144 | *L. thermotolerans* | 1.112 | 1.048 | 55 | 8.4 | 4.12 | 3.09 |
| YH156 | *S. japonicus* | 1.112 | 1.038 | 64 | 9.7 | 4.12 | 3.35 |
| C1 | *S. cerevisiae* | 1.112 | 1.006 | 94 | 13.9 | 4.12 | 3.51 |
| C2 | *S. cerevisiae* | 1.112 | 1.002 | 98 | 14.4 | 4.12 | 3.70 |

<sup>a</sup> Abbreviations: OG, original gravity; FG, final gravity; ABV, alcohol by volume.

Broadly comparing the LAY-fermented meads to the controls, it is obvious that as a group, they were significantly less attenuated and, consequently, had lower ABVs than meads produced with WLP715 (*p* = 0.0003; Fig. 1A,B). This was not unexpected because WLP715 is an industrial strain known for its high levels of attenuation (15), while the LAY have displayed variable levels of attenuation of beer wort (14) and mead (17). Although all 15 meads had a final pH of 3.70 or below, those made with LAY were significantly more acidic on average (*p* = 0.0004; Fig. 1C). This is consistent with their heterofermentative nature and ability to sour beer by lactic acidification (14).

**Figure 1.**
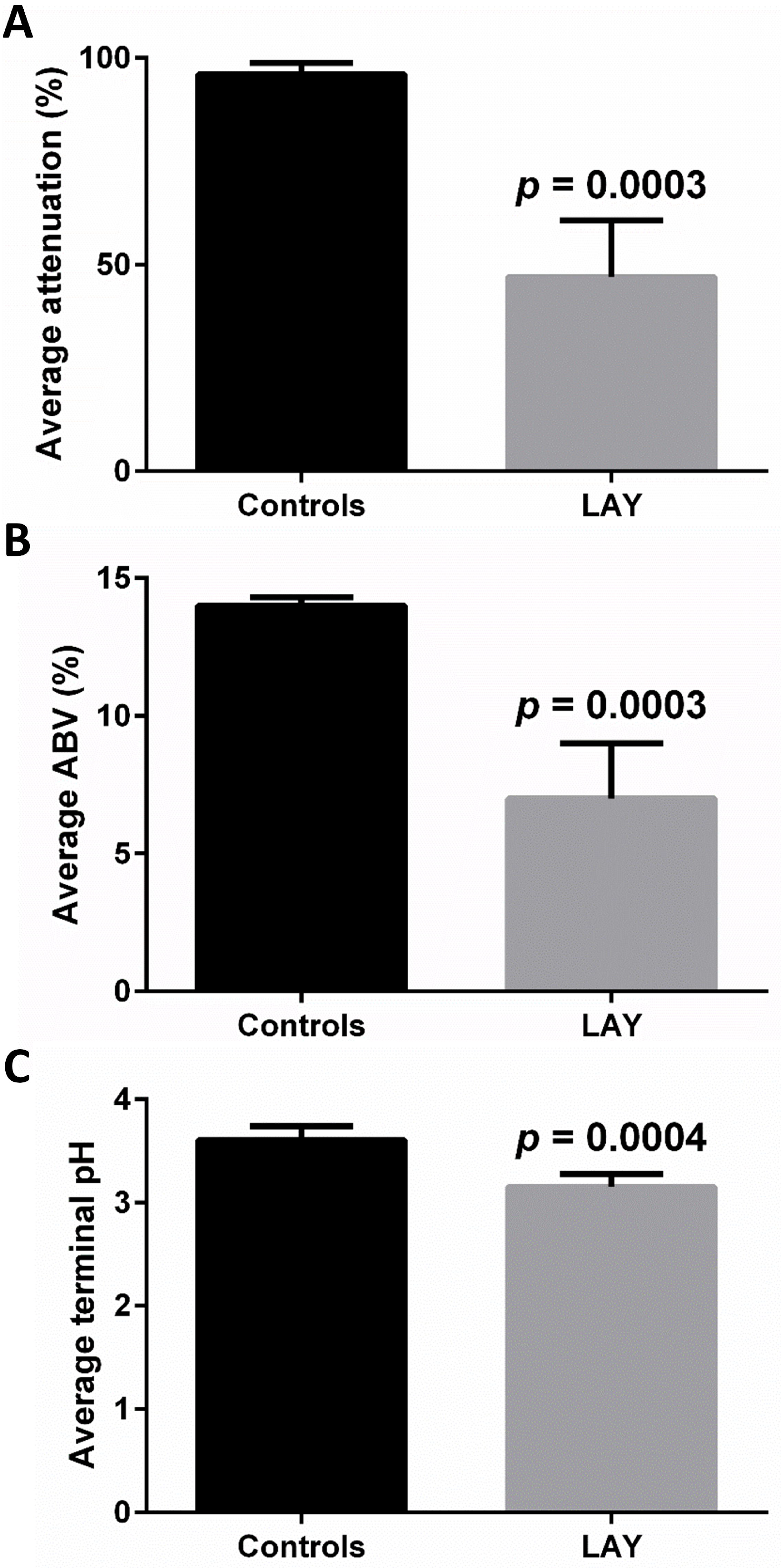
Average attenuation, ABV, and terminal pH of the experimental meads. Comparisons between the mean attenuation, ABV, and pH of the meads produced in the control fermentations (black) and LAY fermentations (gray). The error bars correspond to the standard deviations (SD). Values were compared by *t* tests using the Holm-Sidak method, with a = 5% and without assuming a consistent SD. Significant *p*-values are listed on the graphs.

### 3.2. Sensory analysis

Various methods have been used for the sensory analysis of mead (*e.g.*, (18,19)). At its root, sensory analysis is subjective, being influenced by multiple factors including the experience level of the participants and various methods of analysis (20). To simulate consumer feedback, participants in our sensory analysis were all mead enthusiasts but varied from self-described novices to experts. We received and documented the results recorded on 490 sensory analysis feedback sheets. The overall impression scores calculated from the feedback (see Section 2.4.) are shown in Figure 2. Eight of the LAY strains yielded meads that scored better than the average of 86 ± 7, with the YH77 mead scoring highest at 107 points. The mead fermented by YH26 scored lowest at 56. Further trends in aroma and flavor perception are detailed below.

**Figure 2.**
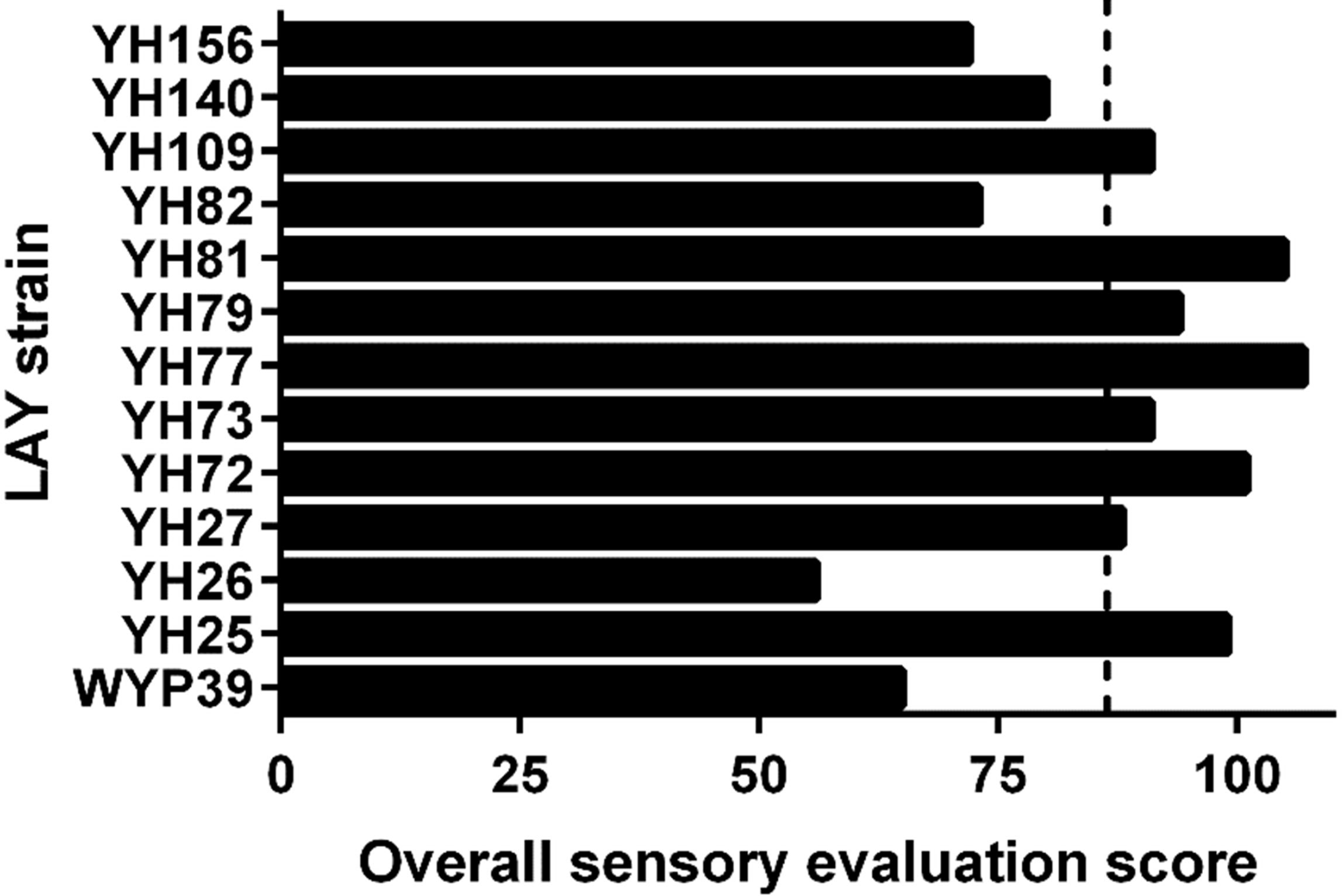
Overall impression scores for the LAY meads. Each sensory analyst gave the LAY meads an overall impression ranking from 1 (least favorable) to 5 (most favorable), and those scores are tallied here. The dashed line indicates the average overall impression score of the 13 experimental meads.

#### 3.2.1. Aroma

Next, we calculated the average general aroma rating for a series of descriptors: delicate, light, mild, pleasant, pungent, rich, sharp, and strong. No two meads had identical profiles, though several were quite similar (*e.g.*, YH73 and YH77 meads) to the average of the group, which tended toward light and mild (Fig. 3). Overall, the aromas were favorable, though several LAY produced meads whose aromatics stood out as being pungent (YH26 and YH156) and/or sharp (YH79 and YH156).

**Figure 3.**
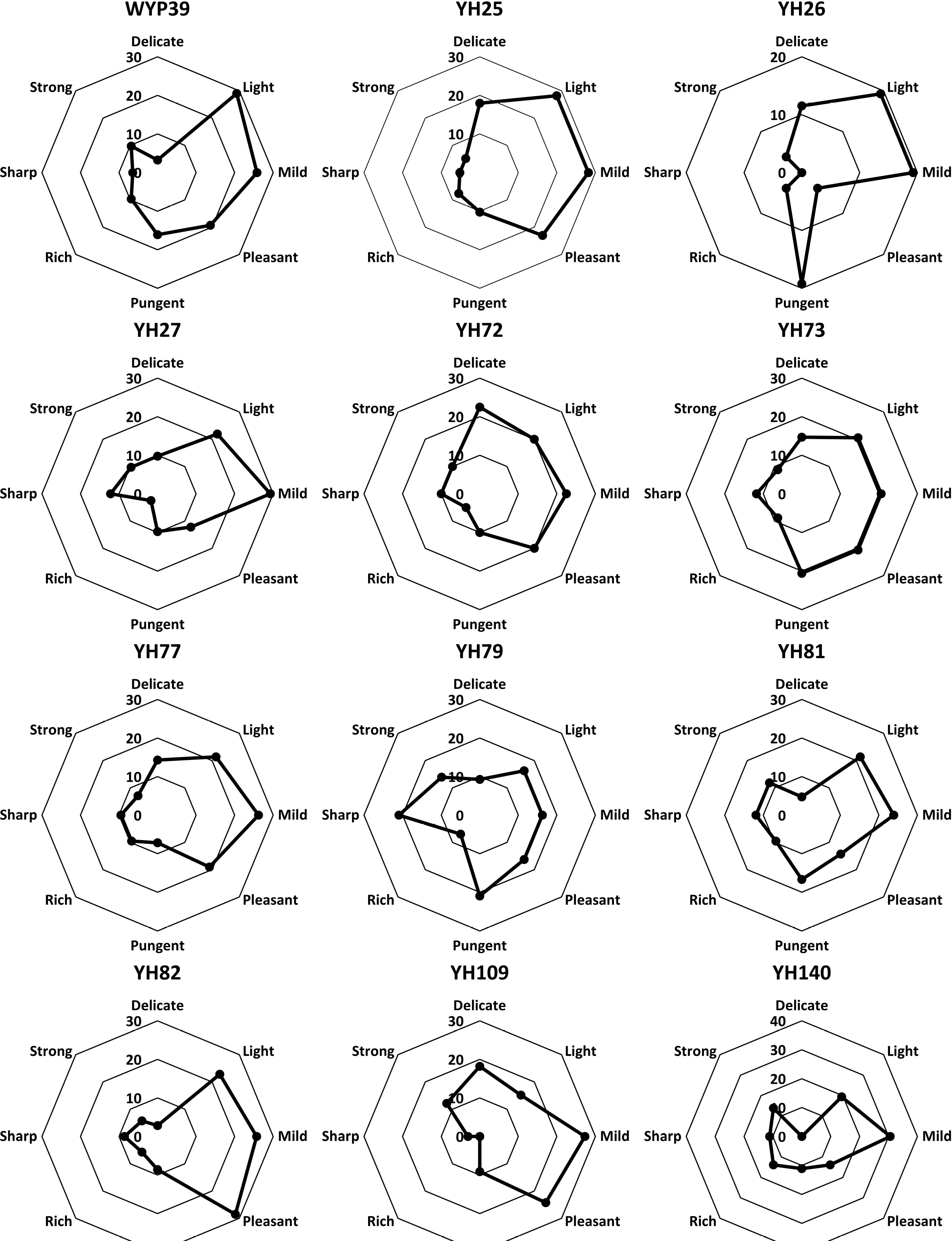
General aroma ratings. The average scores for the indicated descriptors were calculated and plotted as radar charts. Not every survey participant sampled every mead, so averages were calculated from between 26 and 43 evaluations for each mead depending on the number of tasters. The LAY strain used to ferment the mead is indicated above the plotted data. The “Average” label indicates that the data are the means across all 13 experimental meads. Mild, light, and pleasant aromas dominated the meads.

The sensory analysts also scored aromatic esters with the following descriptors: apple, banana, berry, citrus, clean, floral, fruity, grape, mango, melon, nail polish, pear, ripe fruit, stone fruit, tropical, and unripe fruit. On average, the meads displayed citrus and fruity notes (Fig. 4). Among the 13 experimental meads, the results were more mixed here than above. Some were ranked highly for banana aromatics (YH27 and YH156), while others had stronger tropical notes (YH25 and YH77) or were more variably scored (WYP39 and YH82).

**Figure 4.**
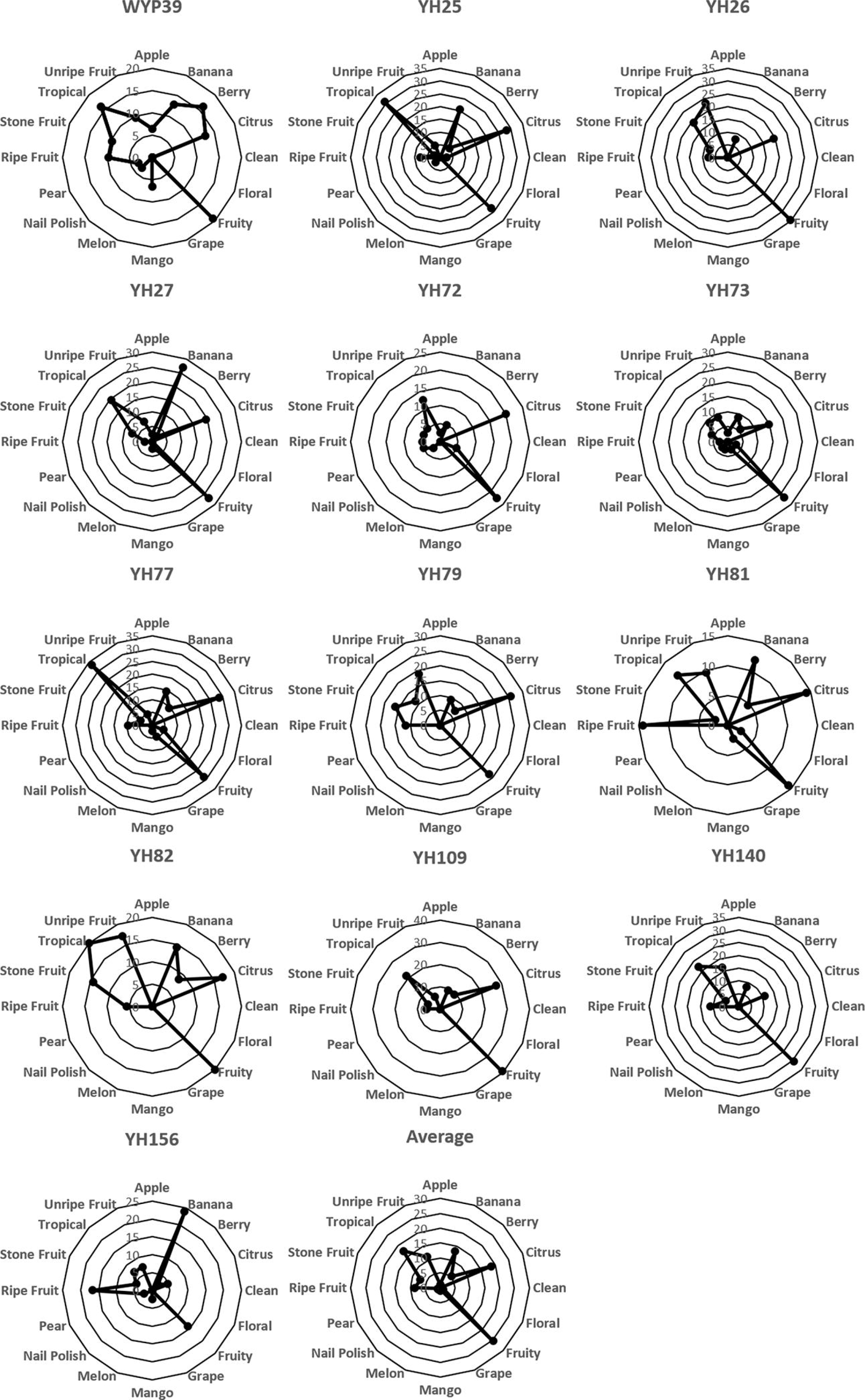
Aromatic ester ratings. The average scores for the indicated aroma descriptors were calculated and plotted as described in Figure 3. The aromas were predominantly described as citrus, fruity, and tropical.

Finally, “other” aromas were scored for odors reminiscent of acetone, acidic, alcohol, bitter, bread, bubble gum, buttery, candy, cinnamon, earthy, grassy, medicinal, oaky, phenolic, smoky, sour, vegetal, vinous, and yeast descriptors. The average other aroma profile was not dominated by any of the descriptors, highlighting the variability among the experimental meads (Fig. 5). Two meads were reported as highly buttery (YH26 and YH77), two were earthy (YH82 and YH140), and YH156 yielded a mead with a strong acetone aroma.

**Figure 5.**
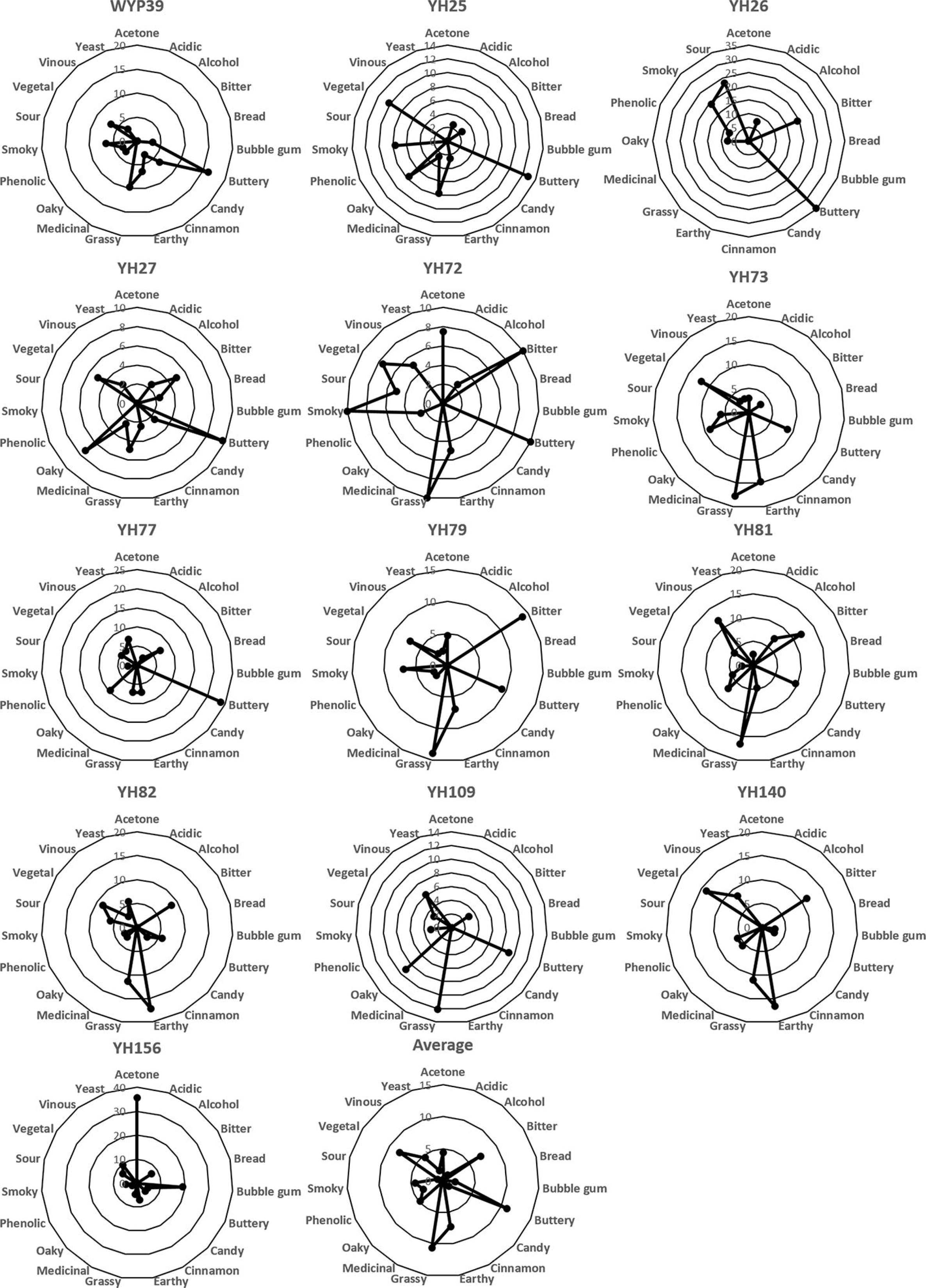
“Other” aroma ratings. The average scores for the indicated aromatic descriptors were calculated and plotted as described in Figure 3. All meads smelled acidic.

#### 3.2.2. Flavor

As above for aroma, we also calculated the average general flavor ratings for the experimental meads using a series of descriptors: bright, delicate, light, mild, pleasant, pungent, rich, sharp, strong, and yeasty. The mean flavors were overall mild and pleasant, but there was some diversity among individual meads (Fig. 6). For instance, the meads produced by YH27, YH81, YH109, and YH156 tasted quite sharp, while the YH72 mead had a rich flavor.

**Figure 6.**
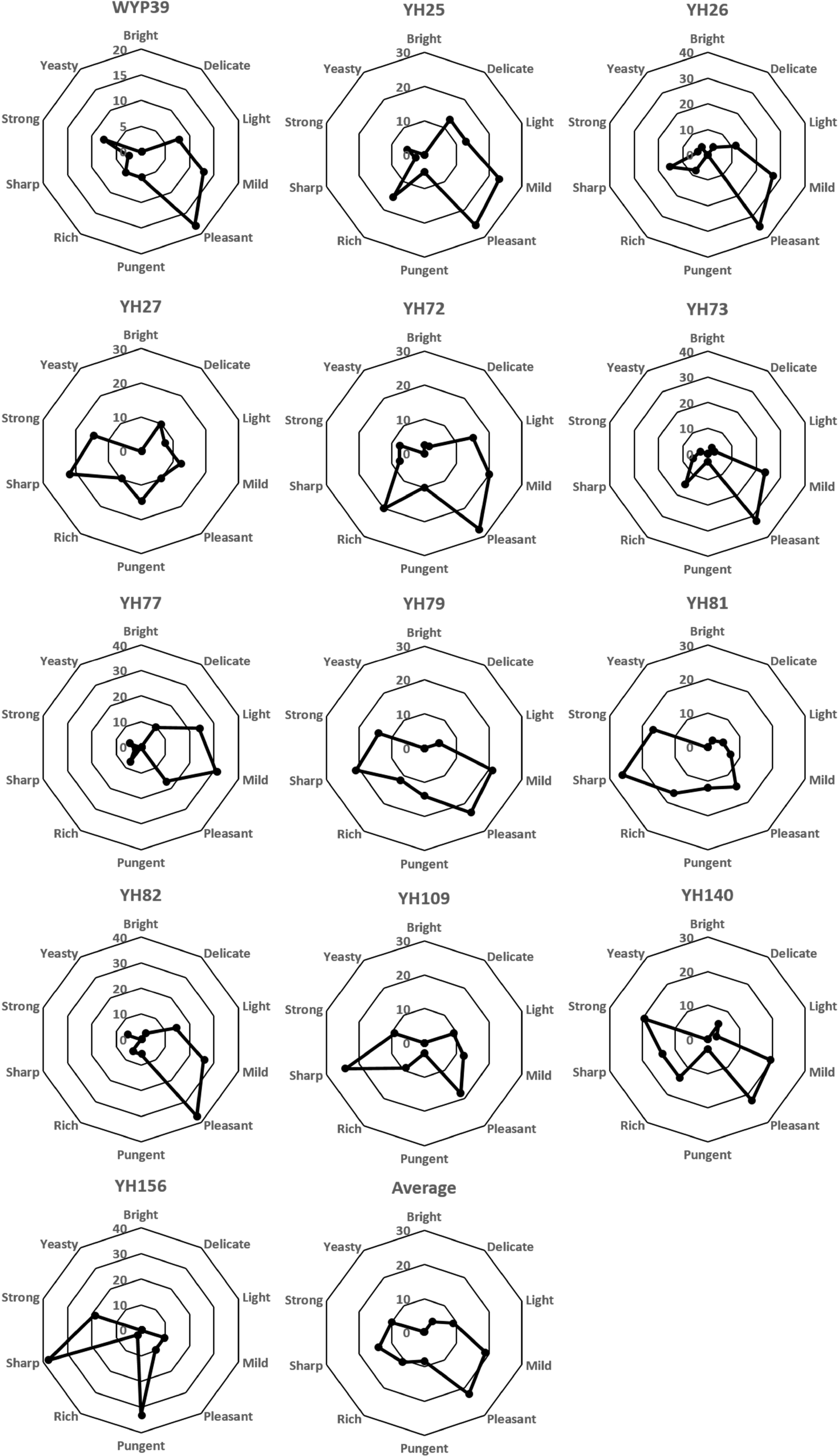
General flavor ratings. The average scores for the indicated flavor descriptors were calculated and plotted as described in Figure 3. The meads tasted mild and pleasant overall.

Flavors were also scored for apple, banana, berry, Brett, citrus, fruity, pear, stone fruit, and tropical descriptors. Here, Brett refers to “Brett character” or the rustic barnyard flavors characteristic of beverages fermented with common strains of *Brettanomyces* spp. (13,21-23). On average, apple, citrus, fruity, and pear flavors predominated, and several individual meads shared this general profile (YH27, YH73, YH77, and YH82) (Fig. 7). Notably, some meads scored highly for citrus and/or pear flavors, such as those made by YH79, YH81, and YH140.

**Figure 7.**
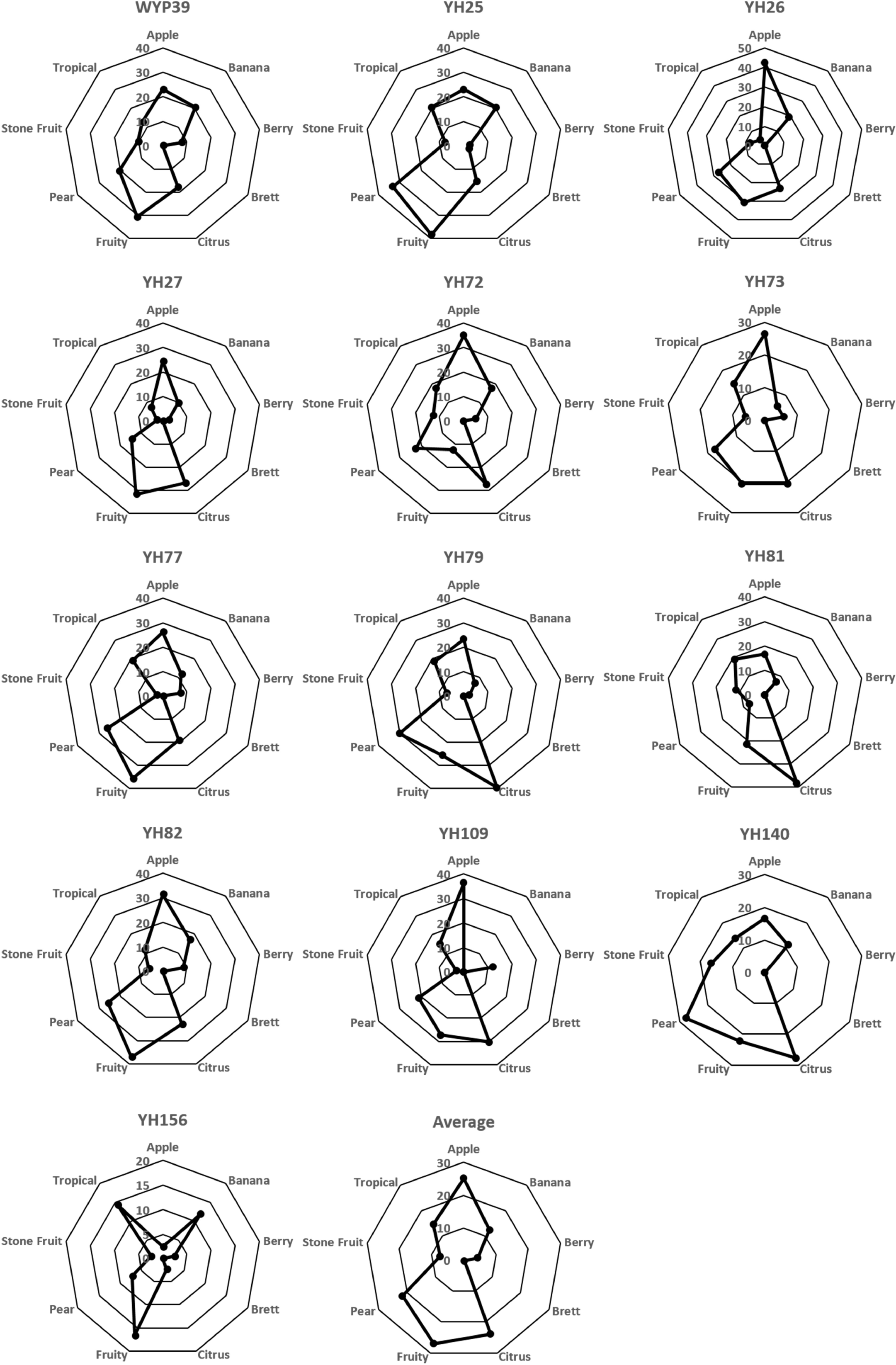
Fruit and Brett flavor ratings. The average scores for fruit-like and Brett character flavor descriptors were calculated and plotted as described in Figure 3. Flavors of apple, citrus, fruit, and pear were common.

Additional flavor descriptors (acetone, astringent, bitter, bread, bubble gum, buttery, earthy, funk, grassy, leather, oaky, silky, smoky, sour, sweet, tart, vegetal, vinous, and yeast) were also considered. Perhaps unsurprisingly, the average profile ranked as highly sour (Fig. 8). Indeed, this was the case for all but two of the individual meads (YH25 and YH77), which were instead perceived as buttery. Several meads were also ranked as bitter: YH27, YH79, and YH156. Concerning the perceived sourness of the meads, however, it should be noted that tasters were informed that the meads were sour, and this may have influenced their feedback. This could explain why the WLP715 control meads were also ranked as having some perceived sour flavor. That said, the terminal pHs of all experimental and control meads were all well below 4.0 (Table 1) due to their organic acid content at the end of fermentation, which corresponds with sour perception (24,25).

**Figure 8.**
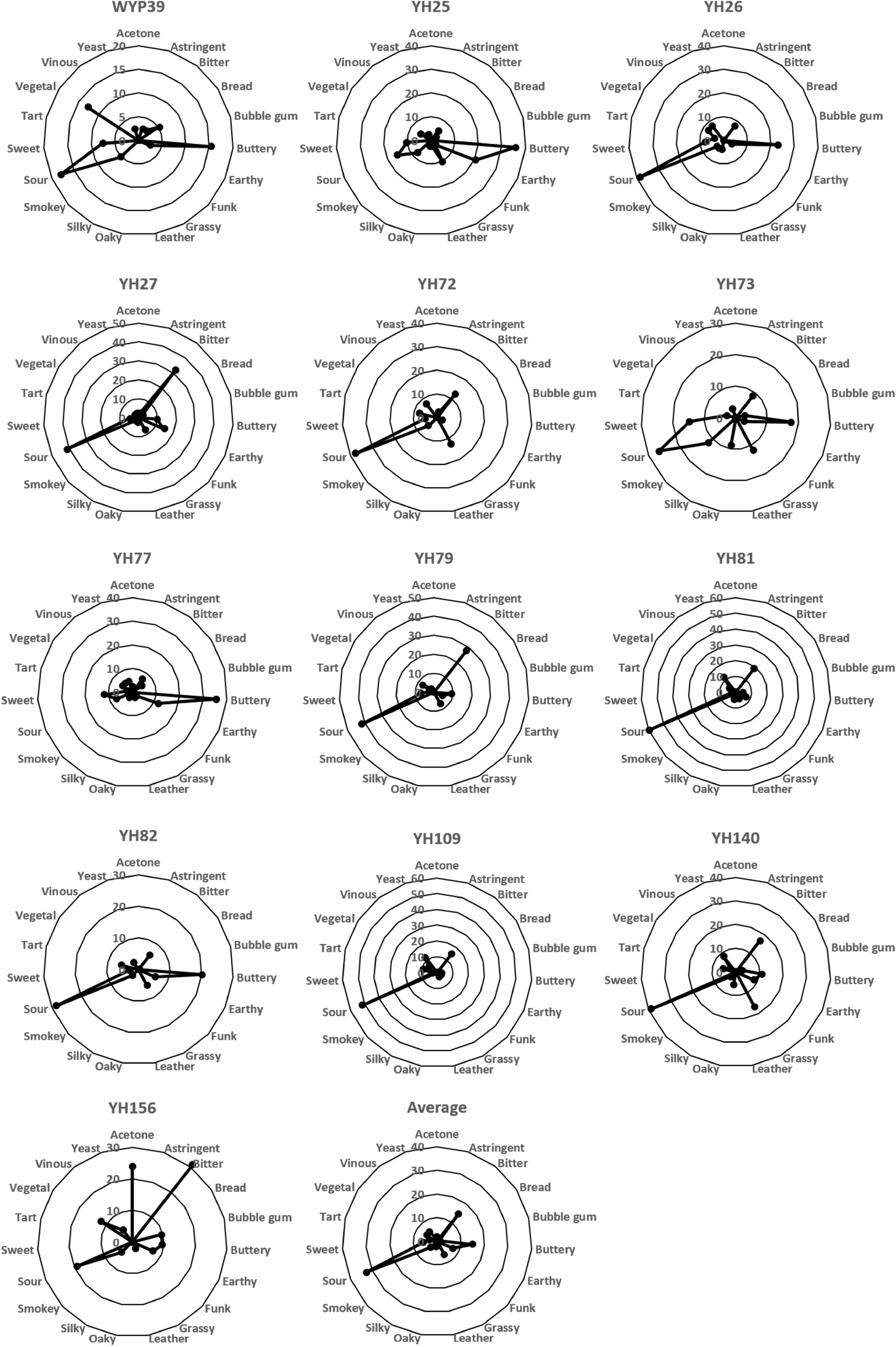
“Other” flavor ratings. The average scores for the indicated flavor descriptors were calculated and plotted as described in Figure 3. All meads scored highly for the sour flavor descriptor.

## Discussion

This project provided insight into the sensory characteristics LAY can provide in mead making. In the overall organoleptic survey of sour meads, most sensory analysts preferred the mead fermented with *L. fermentati strain* YH77, which produced citrusy, tropical aromas with mild, fruity, and pear flavors. In contrast, the least preferred mead was produced by *S. japonicus* strain YH156, which displayed pungent banana and acetone aromas, as well as strong, bitter, and acetone flavors. It is interesting to note that even though YH156 was the least favored strain, it flocculated the most rapidly and exhibited some initially pungent ammonia aromas during fermentation that were not discernible in the finished mead. Regardless, it should be noted that the evaluated sensory data only provide insights into customer preferences and not the absolute quantified intensity of these characteristics. Additional experiments using gas chromatography and mass spectrometry would be needed to compare the concentrations of sensory compounds between meads. One should also be mindful of the effects of other additives on the final sensory attributes of the meads as well. For instance, the choice of fining agents is known to impact the organoleptic profiles of meads (26,27). We did not test such variables in this work, but their influence on the final sour perception of meads fermented by LAY deserves attention.

The LAY strains tested here tended to attenuate mead less than they do beer wort (17,28), with *L. thermotolerans* strain YH27 yielding the highest attenuation, finishing with an ABV of 10.2%. *H. vineae* strain YH72 had the lowest attenuation, with an ABV of 4.2%, which is in stark contrast to our recent work with YH72 in mead production that yielded a finished product with 8.2% ABV (17). The reason for this discrepancy is currently unknown but could be due to variations in the honey musts used between experiments or the yeast nutrient supplementation regime used here. There was also some stochastic variation in the attenuation of the meads (*e.g.*, the FGs of C1 and C2 were 1.006 and 1.002 despite identical must, nutrient, inoculation, and fermentation conditions). Additionally, the terminal pH also varied among the experimental meads, with *L. thermotolerans* strain YH79 producing the lowest at pH = 3.0 and the highest terminal pH being found in meads made by YH77 and YH156 (both finishing at pH = 3.35). It is interesting to note that of the four strains with the lowest terminal pH (YH79, pH = 3.0; YH109, pH = 3.01; YH81 pH = 3.02; and YH72 pH = 3.07), YH81 mead scored as having the highest perceived sour flavors (Fig. 8), and the YH72 and YH82 meads had the highest perceived sour aromas (Fig. 5).

The sensory data gathered from this initial study of LAY-fermented meads makes the case for future work with these organisms. For instance, it would be interesting to examine the effects of fermentation temperature on the organoleptic properties of LAY meads, the effects of initial honey must gravity on attenuation, and the alcohol tolerances of these yeast strains. We also suggest the need for the addition of a sour mead category into the existing BJCP Mead Styles to encourage such experimentation by professional and home mead makers using LAY and other more common souring microbes.

Given that the yeast strains used here produced such an array of aromas and flavors in a fundamental semi-sweet traditional mead, paired with the variety of attenuation levels and levels of clarity yielded by traditional souring techniques, the addition of a new sour mead category will help to encompass the LAY attributes. Indeed, a sour mead category would allow such meads produced by a variety of souring methods to be further evaluated with regard to their unique organoleptic characteristics, resulting in potential growth in what is already a popular style among all levels of mead drinkers. Sour mead could potentially win favor with the already established sour beer enthusiasts and encourage kombucha and other fermented product consumers and brewers to try sour meads, thus expanding the already growing mead market.

## Acknowledgements

Funding for this project was provided by a grant from the American Mead Makers Association. We thank all of the sensory analysts for their feedback.

## Abbreviations

LAY: lactic acid-producing yeast

ABV: alcohol by volume

OG: original gravity

FG: final gravity

SG: specific gravity

AMMA: American Mead Makers Association

TOSNA: traditional organic staggered nutrient addition

