## Supplemental materials for "An organoleptic survey of meads made with lactic acid-producing yeasts"

Table S1. Gravity measurements and tasting notes throughout fermentation. (Table S1.xlsx)

Example sensory analysis feedback sheets for strain YH25.

### AMMA Sour Mead Tasting Feed Back Sheet

The purpose of this session is to gather as much feedback as possible on the **FLAVOR** profiles and attributes for the 13 different bacteria free, souring yeast strains used in these meads.

As a taster, do you consider yourself ☐ Novice ☐ Experienced ☐ Seasoned ☐ Professional

Check as many of the boxes for any **Bouquet/Aroma** (Smell) and **Flavor** (Taste) attributes you perceive for the mead.

---

#### SOUR MEAD 1 Yeast: YH25

**BOUQUET/AROMA**

| **Descriptors** | **Esters** | **Other** | **Comments** |
|---|---|---|---|
| ☐ Delicate | ☑ Fruity | ☐ Bitter | |
| ☐ Light | ☐ Banana | ☐ Buttery | |
| ☐ Mild | ☑ Berry | ☐ Earthy | |
| ☑ Pleasant | ☐ Citrus | ☐ Grassy | |
| ☑ Rich | ☑ Tropical | ☐ Oaky | |
| ☐ Strong | ☐ Stone Fruit | ☐ Smoky | |
| ☐ Pungent | ☐ Unripe Fruit | ☐ Vegetal | |
| ☐ Sharp | ☑ Ripe Fruit | ☐ Vinous | |
| ☐ ________ | ☐ ________ | ☐ ________ | |

**FLAVORS**

| **Descriptors** | **Esters** | **Other** | **Comments** |
|---|---|---|---|
| ☐ Delicate | ☑ Apple | ☐ Bitter | |
| ☐ Light | ☐ Banana | ☑ Buttery | |
| ☐ Mild | ☐ Berry | ☑ Earthy | |
| ☐ Pleasant | ☐ Citrus | ☐ Grassy | |
| ☐ Rich | ☐ Fruity | ☐ Oaky | |
| ☐ Strong | ☐ Pear | ☐ Smoky | |
| ☐ Pungent | ☐ Stone Fruit | ☐ Sour | |
| ☐ Sharp | ☐ Tropical | ☐ Vegetal | |
| ☑ yeasty | ☐ ________ | ☐ Vinous | |
| ☐ ________ | ☐ ________ | ☐ ________ | |

If this yeast were commercially available, would you buy it? ☑ No ☐ Maybe ☐ Absolutely

**OVERALL IMPRESSION**

☐ Yuck! ☑ Meh! ☐ Like it! ☐ Love it! ☐ Got to have it!

#### AMMA Sour Mead Tasting Feed Back Sheet

The purpose of this session is to gather as much feedback as possible on the **FLAVOR** profiles and attributes for the 13 different bacteria free, souring yeast strains used in these meads.

As a taster, do you consider yourself ❑ Novice ❑ Experienced ❑ Seasoned ❑ Professional

Check as many of the boxes for any **Bouquet/Aroma** (Smell) and **Flavor** (Taste) attributes you perceive for the mead.

---

#### SOUR MEAD 1 Yeast: YH25

##### BOUQUET/AROMA

| Descriptors | Esters | Other | Comments |
|---|---|---|---|
| ❑ Delicate | ❑ Fruity | ❑ Bitter | |
| ❑ Light | ❑ Banana | ❑ Buttery | |
| ❑ Mild | ❑ Berry | ❑ Earthy | |
| ❑ Pleasant | ❑ Citrus | ❑ Grassy | |
| ❑ Rich | ❑ Tropical | ❑ Oaky | |
| ❑ Strong | ❑ Stone Fruit | ❑ Smoky | |
| ❑ Pungent | ❑ Unripe Fruit | ❑ Vegetal | |
| ❑ Sharp | ❑ Ripe Fruit | ❑ Vinous | |
| ❑ ________ | ❑ ________ | ❑ ________ | |

##### FLAVORS

| Descriptors | Esters | Other | Comments |
|---|---|---|---|
| ❑ Delicate | ❑ Apple | ❑ Bitter | |
| ❑ Light | ❑ Banana | ❑ Buttery | |
| ❑ Mild | ❑ Berry | ❑ Earthy | |
| ❑ Pleasant | ❑ Citrus | ❑ Grassy | |
| ❑ Rich | ❑ Fruity | ❑ Oaky | |
| ❑ Strong | ❑ Pear | ❑ Smoky | |
| ❑ Pungent | ❑ Stone Fruit | ❑ Sour | |
| ❑ Sharp | ❑ Tropical | ❑ Vegetal | |
| ❑ ________ | ❑ ________ | ❑ Vinous | |
| ❑ ________ | ❑ ________ | ❑ ________ | |

If this yeast were commercially available, would you buy it? ❑ No ❑ Maybe ❑ Absolutely

##### OVERALL IMPRESSION

❑ Yuck! ❑ Meh! ❑ Like it! ❑ Love it! ❑ Got to have it!

### AMMA Sour Mead Tasting Feed Back Sheet

The purpose of this session is to gather as much feedback as possible on the **FLAVOR** profiles and attributes for the 13 different bacteria free, souring yeast strains used in these meads.

As a taster, do you consider yourself ☐ Novice ☐ Experienced ☐ Seasoned ☐ Professional

Check as many of the boxes for any **Bouquet/Aroma** (Smell) and **Flavor** (Taste) attributes you perceive for the mead.

---

#### SOUR MEAD 1 Yeast: YH25

##### BOUQUET/AROMA

| Descriptors | Esters | Other | Comments |
|---|---|---|---|
| ☒ Delicate | ☐ Fruity | ☐ Bitter | |
| ☐ Light | ☐ Banana | ☐ Buttery | |
| ☐ Mild | ☒ Berry | ☐ Earthy | |
| ☐ Pleasant | ☐ Citrus | ☒ Grassy | |
| ☐ Rich | ☐ Tropical | ☐ Oaky | |
| ☐ Strong | ☐ Stone Fruit | ☐ Smoky | |
| ☐ Pungent | ☐ Unripe Fruit | ☐ Vegetal | |
| ☐ Sharp | ☐ Ripe Fruit | ☐ Vinous | |
| ☐ ________ | ☐ ________ | ☐ ________ | |

##### FLAVORS

| Descriptors | Esters | Other | Comments |
|---|---|---|---|
| ☐ Delicate | ☒ Apple | ☒ Bitter | Very lite bitter edge |
| ☐ Light | ☐ Banana | ☒ Buttery | Maybe a bit "sour" |
| ☒ Mild | ☐ Berry | ☐ Earthy | |
| ☐ Pleasant | ☐ Citrus | ☐ Grassy | |
| ☐ Rich | ☐ Fruity | ☐ Oaky | |
| ☐ Strong | ☒ Pear | ☐ Smoky | |
| ☐ Pungent | ☐ Stone Fruit | ☐ Sour | |
| ☐ Sharp | ☐ Tropical | ☐ Vegetal | |
| ☐ ________ | ☐ ________ | ☐ Vinous | |
| ☐ ________ | ☐ ________ | ☐ ________ | |

If this yeast were commercially available, would you buy it? ☒ No ☐ Maybe ☐ Absolutely

##### OVERALL IMPRESSION

☐ Yuck! ☐ Meh! X ☐ Like it! ☐ Love it! ☐ Got to have it!

### AMMA Sour Mead Tasting Feed Back Sheet

The purpose of this session is to gather as much feedback as possible on the **FLAVOR** profiles and attributes for the 13 different bacteria free, souring yeast strains used in these meads.

As a taster, do you consider yourself ☐ Novice ☐ Experienced ☑ Seasoned ☐ Professional

Check as many of the boxes for any **Bouquet/Aroma** (Smell) and **Flavor** (Taste) attributes you perceive for the mead.

---

#### SOUR MEAD 1 Yeast: YH25

##### BOUQUET/AROMA

| Descriptors | Esters | Other | Comments |
|---|---|---|---|
| ☐ Delicate | ☐ Fruity | ☐ Bitter | |
| ☐ Light | ☑ Banana | ☐ Buttery | |
| ☑ Mild | ☐ Berry | ☐ Earthy | |
| ☐ Pleasant | ☐ Citrus | ☐ Grassy | |
| ☐ Rich | ☐ Tropical | ☐ Oaky | |
| ☐ Strong | ☐ Stone Fruit | ☐ Smoky | |
| ☑ Pungent | ☐ Unripe Fruit | ☐ Vegetal | |
| ☐ Sharp | ☐ Ripe Fruit | ☐ Vinous | |
| ☐ ________ | ☐ ________ | ☐ ________ | |

##### FLAVORS

| Descriptors | Esters | Other | Comments |
|---|---|---|---|
| ☐ Delicate | ☑ Apple | ☐ Bitter | Diacytel End Aftertaste |
| ☐ Light | ☑ Banana | ☑ Buttery | |
| ☑ Mild | ☐ Berry | ☑ Earthy | |
| ☐ Pleasant | ☐ Citrus | ☐ Grassy | |
| ☐ Rich | ☑ Fruity | ☐ Oaky | |
| ☐ Strong | ☑ Pear | ☐ Smoky | |
| ☑ Pungent | ☐ Stone Fruit | ☐ Sour | |
| ☐ Sharp | ☐ Tropical | ☐ Vegetal | |
| ☐ ________ | ☐ ________ | ☐ Vinous | |
| ☐ ________ | ☐ ________ | ☐ ________ | |

If this yeast were commercially available, would you buy it? ☐ No ☐ Maybe ☐ Absolutely

##### OVERALL IMPRESSION

☐ Yuck! ☑ Meh! ☐ Like it! ☐ Love it! ☐ Got to have it!

### AMMA Sour Mead Tasting Feed Back Sheet

The purpose of this session is to gather as much feedback as possible on the **FLAVOR** profiles and attributes for the 13 different bacteria free, souring yeast strains used in these meads.

As a taster, do you consider yourself ☐ Novice ☐ Experienced ☐ Seasoned ☒ Professional

Check as many of the boxes for any **Bouquet/Aroma** (Smell) and **Flavor** (Taste) attributes you perceive for the mead.

---

#### SOUR MEAD 1 Yeast: YH25

##### BOUQUET/AROMA

| Descriptors | Esters | Other |
|---|---|---|
| ☐ Delicate | ☐ Fruity | ☐ Bitter |
| ☐ Light | ☐ Banana | ☐ Buttery |
| ☐ Mild | ☐ Berry | ☐ Earthy |
| ☐ Pleasant | ☒ Citrus | ☐ Grassy |
| ☒ Rich | ☐ Tropical | ☒ Oaky |
| ☐ Strong | ☐ Stone Fruit | ☐ Smoky |
| ☐ Pungent | ☐ Unripe Fruit | ☐ Vegetal |
| ☐ Sharp | ☐ Ripe Fruit | ☐ Vinous |
| ☐ ________ | ☐ ________ | ☐ ________ |

**Comments**

This was the best entry after checking all

##### FLAVORS

| Descriptors | Esters | Other |
|---|---|---|
| ☐ Delicate | ☐ Apple | ☐ Bitter |
| ☒ Light | ☐ Banana | ☐ Buttery |
| ☐ Mild | ☐ Berry | ☐ Earthy |
| ☐ Pleasant | ☐ Citrus | ☐ Grassy |
| ☐ Rich | ☒ Fruity | ☐ Oaky |
| ☐ Strong | ☐ Pear | ☐ Smoky |
| ☐ Pungent | ☐ Stone Fruit | ☐ Sour |
| ☐ Sharp | ☐ Tropical | ☐ Vegetal |
| ☐ ________ | ☐ ________ | ☒ Vinous |
| ☐ ________ | ☐ ________ | ☐ ________ |

**Comments**

If this yeast were commercially available, would you buy it? ☐ No ☐ Maybe ☒ Absolutely

**OVERALL IMPRESSION**

☐ Yuck! ☐ Meh! ☐ Like it! ☐ Love it! ☒ Got to have it!

#### AMMA Sour Mead Tasting Feed Back Sheet

The purpose of this session is to gather as much feedback as possible on the **FLAVOR** profiles and attributes for the 13 different bacteria free, souring yeast strains used in these meads.

As a taster, do you consider yourself ☐ Novice ☒ Experienced ☐ Seasoned ☐ Professional

Check as many of the boxes for any **Bouquet/Aroma** (Smell) and **Flavor** (Taste) attributes you perceive for the mead.

---

#### SOUR MEAD 1 Yeast: YH25

##### BOUQUET/AROMA

| Descriptors | Esters | Other | Comments |
|---|---|---|---|
| ☐ Delicate | ☐ Fruity | ☐ Bitter | |
| ☐ Light | ☐ Banana | ☐ Buttery | |
| ☐ Mild | ☐ Berry | ☐ Earthy | |
| ☐ Pleasant | ☐ Citrus | ☐ Grassy | |
| ☐ Rich | ☐ Tropical | ☐ Oaky | |
| ☐ Strong | ☐ Stone Fruit | ☐ Smoky | |
| ☐ Pungent | ☐ Unripe Fruit | ☐ Vegetal | |
| ☐ Sharp | ☐ Ripe Fruit | ☐ Vinous | |
| ☐ ______ | ☒ Nail Polish | ☐ ______ | |

##### FLAVORS

| Descriptors | Esters | Other | Comments |
|---|---|---|---|
| ☐ Delicate | ☐ Apple | ☐ Bitter | |
| ☐ Light | ☒ Banana | ☒ Buttery | |
| ☐ Mild | ☐ Berry | ☐ Earthy | |
| ☐ Pleasant | ☐ Citrus | ☐ Grassy | |
| ☐ Rich | ☐ Fruity | ☐ Oaky | |
| ☐ Strong | ☐ Pear | ☐ Smoky | |
| ☐ Pungent | ☐ Stone Fruit | ☐ Sour | |
| ☐ Sharp | ☒ Tropical | ☐ Vegetal | |
| ☐ ______ | ☐ ______ | ☐ Vinous | |
| ☐ ______ | ☐ ______ | ☐ ______ | |

If this yeast were commercially available, would you buy it? ☒ No ☐ Maybe ☐ Absolutely

##### OVERALL IMPRESSION

☐ Yuck! ☒ Meh! ☐ Like it! ☐ Love it! ☐ Got to have it!

### AMMA Sour Mead Tasting Feed Back Sheet

The purpose of this session is to gather as much feedback as possible on the **FLAVOR** profiles and attributes for the 13 different bacteria free, souring yeast strains used in these meads.

As a taster, do you consider yourself ☐ Novice ☒ Experienced ☐ Seasoned ☐ Professional

Check as many of the boxes for any **Bouquet/Aroma** (Smell) and **Flavor** (Taste) attributes you perceive for the mead.

---

#### SOUR MEAD 1 Yeast: YH25

##### BOUQUET/AROMA

| Descriptors | Esters | Other | Comments |
|---|---|---|---|
| ☐ Delicate | ☐ Fruity | ☐ Bitter | Clove |
| ☐ Light | ☐ Banana | ☐ Buttery | |
| ☐ Mild | ☐ Berry | ☐ Earthy | |
| ☐ Pleasant | ☐ Citrus | ☐ Grassy | |
| ☐ Rich | ☐ Tropical | ☐ Oaky | |
| ☐ Strong | ☐ Stone Fruit | ☐ Smoky | |
| ☐ Pungent | ☐ Unripe Fruit | ☐ Vegetal | |
| ☐ Sharp | ☐ Ripe Fruit | ☐ Vinous | |
| ☐ neutral | ☐ ________ | ☐ ________ | |

##### FLAVORS

| Descriptors | Esters | Other | Comments |
|---|---|---|---|
| ☐ Delicate | ☐ Apple | ☐ Bitter | |
| ☐ Light | ☐ Banana | ☒ Buttery | |
| ☐ Mild | ☐ Berry | ☐ Earthy | |
| ☐ Pleasant | ☐ Citrus | ☐ Grassy | |
| ☒ Rich | ☒ Fruity | ☐ Oaky | |
| ☐ Strong | ☐ Pear | ☐ Smoky | |
| ☐ Pungent | ☒ Stone Fruit | ☐ Sour | |
| ☐ Sharp | ☐ Tropical | ☐ Vegetal | |
| ☐ ________ | ☐ ________ | ☐ Vinous | |
| ☐ ________ | ☐ ________ | ☐ ________ | |

If this yeast were commercially available, would you buy it? ☐ No ☒ Maybe ☐ Absolutely

##### OVERALL IMPRESSION

☐ Yuck! ☐ Meh! ☒ Like it! ☐ Love it! ☐ Got to have it!

### AMMA Sour Mead Tasting Feed Back Sheet

The purpose of this session is to gather as much feedback as possible on the **FLAVOR** profiles and attributes for the 13 different bacteria free, souring yeast strains used in these meads.

As a taster, do you consider yourself ☐ Novice ☒ Experienced ☐ Seasoned ☐ Professional

Check as many of the boxes for any **Bouquet/Aroma** (Smell) and **Flavor** (Taste) attributes you perceive for the mead.

---

#### SOUR MEAD 1 Yeast: YH25

##### BOUQUET/AROMA

| Descriptors | Esters | Other | Comments |
|---|---|---|---|
| ☐ Delicate | ☐ Fruity | ☐ Bitter | pleasant |
| ☐ Light | ☒ Banana | ☐ Buttery | |
| ☐ Mild | ☐ Berry | ☐ Earthy | |
| ☐ Pleasant | ☒ Citrus | ☐ Grassy | |
| ☐ Rich | ☐ Tropical | ☐ Oaky | |
| ☐ Strong | ☐ Stone Fruit | ☐ Smoky | |
| ☐ Pungent | ☐ Unripe Fruit | ☐ Vegetal | |
| ☐ Sharp | ☐ Ripe Fruit | ☐ Vinous | |
| ☐ ________ | ☐ ________ | ☐ ________ | |

##### FLAVORS

| Descriptors | Esters | Other | Comments |
|---|---|---|---|
| ☐ Delicate | ☐ Apple | ☐ Bitter | Sweet |
| ☐ Light | ☐ Banana | ☐ Buttery | |
| ☐ Mild | ☐ Berry | ☐ Earthy | |
| ☐ Pleasant | ☐ Citrus | ☐ Grassy | |
| ☒ Rich | ☒ Fruity | ☐ Oaky | |
| ☐ Strong | ☐ Pear | ☐ Smoky | |
| ☐ Pungent | ☐ Stone Fruit | ☐ Sour | |
| ☐ Sharp | ☒ Tropical | ☐ Vegetal | |
| ☐ ________ | ☐ ________ | ☐ Vinous | |
| ☐ ________ | ☐ ________ | ☐ ________ | |

If this yeast were commercially available, would you buy it? ☐ No ☒ Maybe ☐ Absolutely

##### OVERALL IMPRESSION

☐ Yuck! ☐ Meh! ☒ Like it! ☐ Love it! ☐ Got to have it!

### AMMA Sour Mead Tasting Feed Back Sheet

The purpose of this session is to gather as much feedback as possible on the **FLAVOR** profiles and attributes for the 13 different bacteria free, souring yeast strains used in these meads.

As a taster, do you consider yourself ☐ Novice ☐ Experienced ☐ Seasoned ☐ Professional

Check as many of the boxes for any **Bouquet/Aroma** (Smell) and **Flavor** (Taste) attributes you perceive for the mead.

---

#### SOUR MEAD 1 Yeast: YH25

##### BOUQUET/AROMA

| Descriptors | Esters | Other | Comments |
|---|---|---|---|
| ☐ Delicate | ☐ Fruity | ☒ Bitter | |
| ☐ Light | ☒ Banana | ☐ Buttery | |
| ☐ Mild | ☐ Berry | ☐ Earthy | |
| ☐ Pleasant | ☐ Citrus | ☐ Grassy | |
| ☐ Rich | ☐ Tropical | ☐ Oaky | |
| ☐ Strong | ☐ Stone Fruit | ☐ Smoky | |
| ☒ Pungent | ☐ Unripe Fruit | ☐ Vegetal | |
| ☐ Sharp | ☐ Ripe Fruit | ☐ Vinous | |
| ☐ ________ | ☐ ________ | ☐ ________ | |

##### FLAVORS

| Descriptors | Esters | Other | Comments |
|---|---|---|---|
| ☐ Delicate | ☒ Apple | ☐ Bitter | |
| ☐ Light | ☒ Banana | ☒ Buttery | |
| ☒ Mild | ☐ Berry | ☐ Earthy | |
| ☐ Pleasant | ☐ Citrus | ☐ Grassy | |
| ☐ Rich | ☐ Fruity | ☐ Oaky | |
| ☐ Strong | ☐ Pear | ☐ Smoky | |
| ☐ Pungent | ☐ Stone Fruit | ☐ Sour | |
| ☐ Sharp | ☐ Tropical | ☐ Vegetal | |
| ☐ ________ | ☐ ________ | ☐ Vinous | |
| ☐ ________ | ☐ ________ | ☐ ________ | |

If this yeast were commercially available, would you buy it? ☐ No ☒ Maybe ☐ Absolutely

##### OVERALL IMPRESSION

☐ Yuck! ☐ Meh! ☒ Like it! ☐ Love it! ☐ Got to have it!

### AMMA Sour Mead Tasting Feed Back Sheet

The purpose of this session is to gather as much feedback as possible on the **FLAVOR** profiles and attributes for the 13 different bacteria free, souring yeast strains used in these meads.

As a taster, do you consider yourself ☐ Novice ☒ Experienced ☐ Seasoned ☐ Professional

Check as many of the boxes for any **Bouquet/Aroma** (Smell) and **Flavor** (Taste) attributes you perceive for the mead.

---

#### SOUR MEAD 1 Yeast: YH25 RED OAK BARK 6.5%

**BOUQUET/AROMA**

| Descriptors | Esters | Other | Comments |
|---|---|---|---|
| ☐ Delicate | ☐ Fruity | ☐ Bitter | |
| ☐ Light | ☒ Banana | ☐ Buttery | |
| ☐ Mild | ☐ Berry | ☐ Earthy | |
| ☐ Pleasant | ☐ Citrus | ☐ Grassy | |
| ☐ Rich | ☐ Tropical | ☐ Oaky | |
| ☒ Strong | ☐ Stone Fruit | ☐ Smoky | |
| ☒ Pungent | ☐ Unripe Fruit | ☐ Vegetal | |
| ☐ Sharp | ☐ Ripe Fruit | ☐ Vinous | |
| ☐ ________ | ☐ ________ | ☐ ________ | |

**FLAVORS**

| Descriptors | Esters | Other | Comments |
|---|---|---|---|
| ☐ Delicate | ☐ Apple | ☐ Bitter | |
| ☐ Light | ☐ Banana | ☐ Buttery | |
| ☐ Mild | ☐ Berry | ☒ Earthy – DIRT | |
| ☐ Pleasant | ☐ Citrus | ☐ Grassy | |
| ☐ Rich | ☐ Fruity | ☐ Oaky | |
| ☐ Strong | ☐ Pear | ☐ Smoky | |
| ☐ Pungent | ☐ Stone Fruit | ☐ Sour | |
| ☐ Sharp | ☐ Tropical | ☐ Vegetal | |
| ☐ ________ | ☐ ________ | ☐ Vinous | |
| ☐ ________ | ☐ ________ | ☐ ________ | |

If this yeast were commercially available, would you buy it? ☒ No ☐ Maybe ☐ Absolutely

**OVERALL IMPRESSION**

☐ Yuck! ☒ Meh! ☐ Like it! ☐ Love it! ☐ Got to have it!

### AMMA Sour Mead Tasting Feed Back Sheet

The purpose of this session is to gather as much feedback as possible on the **FLAVOR** profiles and attributes for the 13 different bacteria free, souring yeast strains used in these meads.

As a taster, do you consider yourself ☐ Novice ☐ Experienced ☐ Seasoned ☐ Professional

Check as many of the boxes for any **Bouquet/Aroma** (Smell) and **Flavor** (Taste) attributes you perceive for the mead.

---

#### SOUR MEAD 1 Yeast: YH25

6.5%

##### BOUQUET/AROMA

| Descriptors | Esters | Other | Comments |
|---|---|---|---|
| ☐ Delicate | ☐ Fruity | ☐ Bitter | Not a fan |
| ☐ Light | ☐ Banana | ☐ Buttery | aroma is clinical |
| ☐ Mild | ☐ Berry | ☐ Earthy | |
| ☐ Pleasant | ☐ Citrus | ☐ Grassy | |
| ☐ Rich | ☐ Tropical | ☐ Oaky | |
| ☒ Strong | ☐ Stone Fruit | ☐ Smoky | |
| ☒ Pungent | ☐ Unripe Fruit | ☐ Vegetal | |
| ☒ Sharp | ☐ Ripe Fruit | ☐ Vinous | |
| ☐ ______ | ☐ ______ | ☒ Medicinal | |

##### FLAVORS

| Descriptors | Esters | Other | Comments |
|---|---|---|---|
| ☐ Delicate | ☐ Apple | ☐ Bitter | |
| ☐ Light | ☐ Banana | ☐ Buttery | |
| ☐ Mild | ☐ Berry | ☒ Earthy | |
| ☐ Pleasant | ☐ Citrus | ☒ Grassy | |
| ☐ Rich | ☐ Fruity | ☐ Oaky | |
| ☐ Strong | ☐ Pear | ☐ Smoky | |
| ☐ Pungent | ☐ Stone Fruit | ☐ Sour | |
| ☐ Sharp | ☐ Tropical | ☐ Vegetal | |
| ☐ ______ | ☐ ______ | ☐ Vinous | |
| ☐ ______ | ☐ ______ | ☐ ______ | |

If this yeast were commercially available, would you buy it? ☒ No ☐ Maybe ☐ Absolutely

##### OVERALL IMPRESSION

☒ Yuck! ☐ Meh! ☐ Like it! ☐ Love it! ☐ Got to have it!

#### AMMA Sour Mead Tasting Feed Back Sheet

The purpose of this session is to gather as much feedback as possible on the **FLAVOR** profiles and attributes for the 13 different bacteria free, souring yeast strains used in these meads.

As a taster, do you consider yourself ☒ Novice ☐ Experienced ☐ Seasoned ☐ Professional

Check as many of the boxes for any **Bouquet/Aroma** (Smell) and **Flavor** (Taste) attributes you perceive for the mead.

---

#### SOUR MEAD 1 Yeast: YH25

##### BOUQUET/AROMA

| Descriptors | Esters | Other | Comments |
|---|---|---|---|
| ☐ Delicate | ☒ Fruity | ☐ Bitter | |
| ☒ Light | ☐ Banana | ☐ Buttery | |
| ☒ Mild | ☐ Berry | ☐ Earthy | |
| ☐ Pleasant | ☐ Citrus | ☒ Grassy | |
| ☐ Rich | ☐ Tropical | ☐ Oaky | |
| ☐ Strong | ☐ Stone Fruit | ☐ Smoky | |
| ☐ Pungent | ☐ Unripe Fruit | ☐ Vegetal | |
| ☐ Sharp | ☐ Ripe Fruit | ☐ Vinous | |
| ☐ ________ | ☐ ________ | ☐ ________ | |

##### FLAVORS

| Descriptors | Esters | Other | Comments |
|---|---|---|---|
| ☐ Delicate | ☐ Apple | ☐ Bitter | |
| ☐ Light | ☒ Banana | ☒ Buttery | |
| ☒ Mild | ☐ Berry | ☐ Earthy | |
| ☒ Pleasant | ☐ Citrus | ☐ Grassy | |
| ☐ Rich | ☐ Fruity | ☐ Oaky | |
| ☐ Strong | ☐ Pear | ☐ Smoky | |
| ☐ Pungent | ☐ Stone Fruit | ☒ Sour | |
| ☐ Sharp | ☐ Tropical | ☐ Vegetal | |
| ☐ ________ | ☐ ________ | ☐ Vinous | |
| ☐ ________ | ☐ ________ | ☐ ________ | |

If this yeast were commercially available, would you buy it? ☐ No ☒ Maybe ☐ Absolutely

##### OVERALL IMPRESSION

☐ Yuck! ☐ Meh! ☐ Like it! ☒ Love it! ☐ Got to have it!

### AMMA Sour Mead Tasting Feed Back Sheet

The purpose of this session is to gather as much feedback as possible on the **FLAVOR** profiles and attributes for the 13 different bacteria free, souring yeast strains used in these meads.

As a taster, do you consider yourself ☐ Novice ☐ Experienced ☐ Seasoned ☒ Professional

Check as many of the boxes for any **Bouquet/Aroma** (Smell) and **Flavor** (Taste) attributes you perceive for the mead.

---

#### SOUR MEAD 1 Yeast: YH25

##### BOUQUET/AROMA

| Descriptors | Esters | Other |
|---|---|---|
| ☐ Delicate | ☐ Fruity | ☐ Bitter |
| ☒ Light | ☐ Banana | ☐ Buttery |
| ☒ Mild | ☐ Berry | ☐ Earthy |
| ☐ Pleasant | ☐ Citrus | ☐ Grassy |
| ☐ Rich | ☐ Tropical | ☐ Oaky |
| ☐ Strong | ☐ Stone Fruit | ☒ Smoky |
| ☐ Pungent | ☐ Unripe Fruit | ☐ Vegetal |
| ☐ Sharp | ☐ Ripe Fruit | ☐ Vinous |
| ☐ ______ | ☐ ______ | ☐ ______ |

**Comments**

No nose to speak of, slight smokiness

##### FLAVORS

| Descriptors | Esters | Other |
|---|---|---|
| ☒ Delicate | ☐ Apple | ☒ Bitter |
| ☐ Light | ☐ Banana | ☐ Buttery |
| ☐ Mild | ☐ Berry | ☐ Earthy |
| ☒ Pleasant | ☐ Citrus | ☒ Grassy |
| ☐ Rich | ☐ Fruity | ☐ Oaky |
| ☐ Strong | ☒ Pear | ☒ Smoky |
| ☐ Pungent | ☐ Stone Fruit | ☐ Sour |
| ☐ Sharp | ☐ Tropical | ☐ Vegetal |
| ☐ ______ | ☐ ______ | ☐ Vinous |
| ☐ ______ | ☐ ______ | ☐ ______ |

**Comments**

Complex. Definitely grew on me! Worth further exploration

If this yeast were commercially available, would you buy it? ☐ No ☒ Maybe ☐ Absolutely

**OVERALL IMPRESSION**

☐ Yuck! ☐ Meh! ☒ Like it! ☐ Love it! ☐ Got to have it!

### AMMA Sour Mead Tasting Feed Back Sheet

The purpose of this session is to gather as much feedback as possible on the **FLAVOR** profiles and attributes for the 13 different bacteria free, souring yeast strains used in these meads.

As a taster, do you consider yourself ☐ Novice ☐ Experienced [x] Seasoned ☐ Professional

Check as many of the boxes for any **Bouquet/Aroma** (Smell) and **Flavor** (Taste) attributes you perceive for the mead.

---

#### SOUR MEAD 1 Yeast: YH25

##### BOUQUET/AROMA

| Descriptors | Esters | Other | Comments |
|---|---|---|---|
| ☐ Delicate | ☐ Fruity | ☐ Bitter | |
| ☐ Light | ☐ Banana | ☐ Buttery | |
| ☐ Mild | ☐ Berry | ☐ Earthy | |
| ☐ Pleasant | [x] Citrus | ☐ Grassy | |
| ☐ Rich | [x] Tropical | ☐ Oaky | |
| ☐ Strong | ☐ Stone Fruit | ☐ Smoky | |
| ☐ Pungent | ☐ Unripe Fruit | ☐ Vegetal | |
| ☐ Sharp | ☐ Ripe Fruit | ☐ Vinous | |
| ☐ ________ | ☐ ________ | ☐ ________ | |

##### FLAVORS

| Descriptors | Esters | Other | Comments |
|---|---|---|---|
| ☐ Delicate | ☐ Apple | ☐ Bitter | |
| ☐ Light | ☐ Banana | ☐ Buttery | |
| ☐ Mild | ☐ Berry | ☐ Earthy | |
| ☐ Pleasant | ☐ Citrus | ☐ Grassy | |
| ☐ Rich | ☐ Fruity | ☐ Oaky | |
| ☐ Strong | [x] Pear | ☐ Smoky | |
| ☐ Pungent | [x] Stone Fruit | ☐ Sour | |
| ☐ Sharp | ☐ Tropical | ☐ Vegetal | |
| ☐ ________ | ☐ ________ | ☐ Vinous | |
| ☐ ________ | ☐ ________ | ☐ ________ | |

If this yeast were commercially available, would you buy it? ☐ No [x] Maybe ☐ Absolutely

**OVERALL IMPRESSION**

☐ Yuck! ☐ Meh! [x] Like it! ☐ Love it! ☐ Got to have it!

#### AMMA Sour Mead Tasting Feed Back Sheet

The purpose of this session is to gather as much feedback as possible on the **FLAVOR** profiles and attributes for the 13 different bacteria free, souring yeast strains used in these meads.

As a taster, do you consider yourself ☐ Novice ☐ Experienced ☐ Seasoned ☐ Professional

Check as many of the boxes for any **Bouquet/Aroma** (Smell) and **Flavor** (Taste) attributes you perceive for the mead.

---

#### SOUR MEAD 1 Yeast: YH25

**BOUQUET/AROMA**

| Descriptors | Esters | Other | Comments |
|---|---|---|---|
| ☐ Delicate | ☐ Fruity | ☐ Bitter | |
| ☐ Light | ☐ Banana | ☐ Buttery | |
| ☒ Mild | ☐ Berry | ☐ Earthy | |
| ☐ Pleasant | ☐ Citrus | ☐ Grassy | |
| ☐ Rich | ☐ Tropical | ☐ Oaky | |
| ☐ Strong | ☐ Stone Fruit | ☐ Smoky | |
| ☐ Pungent | ☐ Unripe Fruit | ☐ Vegetal | |
| ☐ Sharp | ☐ Ripe Fruit | ☐ Vinous | |
| ☐ ________ | ☐ ________ | ☐ ________ | |

**FLAVORS**

| Descriptors | Esters | Other | Comments |
|---|---|---|---|
| ☐ Delicate | ☐ Apple | ☐ Bitter | |
| ☐ Light | ☐ Banana | ☐ Buttery | |
| ☐ Mild | ☐ Berry | ☐ Earthy | |
| ☐ Pleasant | ☐ Citrus | ☐ Grassy | |
| ☐ Rich | ☐ Fruity | ☐ Oaky | |
| ☐ Strong | ☐ Pear | ☐ Smoky | |
| ☐ Pungent | ☐ Stone Fruit | ☒ Sour | |
| ☒ Sharp | ☐ Tropical | ☐ Vegetal | |
| ☐ ________ | ☐ ________ | ☐ Vinous | |
| ☐ ________ | ☐ ________ | ☐ ________ | |

If this yeast were commercially available, would you buy it? ☒ No ☐ Maybe ☐ Absolutely

**OVERALL IMPRESSION**

☐ Yuck! ☐ Meh! ☐ Like it! ☐ Love it! ☐ Got to have it!

### AMMA Sour Mead Tasting Feed Back Sheet

The purpose of this session is to gather as much feedback as possible on the **FLAVOR** profiles and attributes for the 13 different bacteria free, souring yeast strains used in these meads.

As a taster, do you consider yourself ☑ Novice ☐ Experienced ☐ Seasoned ☐ Professional

Check as many of the boxes for any **Bouquet/Aroma** (Smell) and **Flavor** (Taste) attributes you perceive for the mead.

---

#### SOUR MEAD 1 Yeast: YH25

**BOUQUET/AROMA**

| **Descriptors** | **Esters** | **Other** | **Comments** |
|---|---|---|---|
| ☐ Delicate | ☐ Fruity | ☐ Bitter | ______ |
| ☑ Light | ☐ Banana | ☐ Buttery | ______ |
| ☐ Mild | ☐ Berry | ☐ Earthy | ______ |
| ☐ Pleasant | ☐ Citrus | ☐ Grassy | ______ |
| ☐ Rich | ☐ Tropical | ☐ Oaky | ______ |
| ☐ Strong | ☐ Stone Fruit | ☐ Smoky | ______ |
| ☐ Pungent | ☐ Unripe Fruit | ☐ Vegetal | ______ |
| ☐ Sharp | ☐ Ripe Fruit | ☐ Vinous | |
| ☐ ______ | ☐ ______ | ☐ ______ | |

**FLAVORS**

| **Descriptors** | **Esters** | **Other** | **Comments** |
|---|---|---|---|
| ☐ Delicate | ☑ Apple | ☐ Bitter | ______ |
| ☐ Light | ☑ Banana | ☑ Buttery | ______ |
| ☑ Mild | ☑ Berry | ☐ Earthy | ______ |
| ☐ Pleasant | ☐ Citrus | ☑ Grassy | ______ |
| ☐ Rich | ☐ Fruity | ☐ Oaky | ______ |
| ☐ Strong | ☐ Pear | ☑ Smoky | ______ |
| ☐ Pungent | ☐ Stone Fruit | ☐ Sour | ______ |
| ☐ Sharp | ☐ Tropical | ☐ Vegetal | ______ |
| ☐ ______ | ☐ ______ | ☐ Vinous | ______ |
| ☐ ______ | ☐ ______ | ☐ ______ | |

If this yeast were commercially available, would you buy it? ☐ No ☐ Maybe ☐ Absolutely

**OVERALL IMPRESSION**

☐ Yuck! ☐ Meh! ☑ Like it! ☐ Love it! ☐ Got to have it!

#### AMMA Sour Mead Tasting Feed Back Sheet

The purpose of this session is to gather as much feedback as possible on the **FLAVOR** profiles and attributes for the 13 different bacteria free, souring yeast strains used in these meads.

As a taster, do you consider yourself ☐ Novice ☐ Experienced ☐ Seasoned ☒ Professional

Check as many of the boxes for any **Bouquet/Aroma** (Smell) and **Flavor** (Taste) attributes you perceive for the mead.

---

#### SOUR MEAD 1 Yeast: YH25

##### BOUQUET/AROMA

| Descriptors | Esters | Other | Comments |
|---|---|---|---|
| ☐ Delicate | ☐ Fruity | ☐ Bitter | |
| ☒ Light | ☐ Banana | ☐ Buttery | |
| ☒ Mild | ☐ Berry | ☐ Earthy | |
| ☐ Pleasant | ☒ Citrus | ☐ Grassy | |
| ☐ Rich | ☒ Tropical | ☐ Oaky | |
| ☐ Strong | ☐ Stone Fruit | ☐ Smoky | |
| ☐ Pungent | ☐ Unripe Fruit | ☐ Vegetal | |
| ☐ Sharp | ☐ Ripe Fruit | ☐ Vinous | |
| ☐ ______ | ☐ ______ | ☐ ______ | |

##### FLAVORS

| Descriptors | Esters | Other | Comments |
|---|---|---|---|
| ☐ Delicate | ☐ Apple | ☐ Bitter | Starts fine, but funky aftertaste I don't like |
| ☒ Light | ☐ Banana | ☐ Buttery | |
| ☒ Mild | ☐ Berry | ☐ Earthy | |
| ☒ Pleasant | ☒ Citrus | ☐ Grassy | |
| ☐ Rich | ☒ Fruity | ☐ Oaky | |
| ☐ Strong | ☐ Pear | ☐ Smoky | |
| ☐ Pungent | ☐ Stone Fruit | ☐ Sour | |
| ☐ Sharp | ☐ Tropical | ☐ Vegetal | |
| ☐ ______ | ☐ ______ | ☐ Vinous | |
| ☐ ______ | ☐ ______ | ☒ Funk | |

If this yeast were commercially available, would you buy it? ☐ No ☒ Maybe ☐ Absolutely

##### OVERALL IMPRESSION

☐ Yuck! ☒ Meh! ☐ Like it! ☐ Love it! ☐ Got to have it!

#### AMMA Sour Mead Tasting Feed Back Sheet

The purpose of this session is to gather as much feedback as possible on the **FLAVOR** profiles and attributes for the 13 different bacteria free, souring yeast strains used in these meads.

As a taster, do you consider yourself ☐ Novice ☐ Experienced ☐ Seasoned ☑ Professional

Check as many of the boxes for any **Bouquet/Aroma** (Smell) and **Flavor** (Taste) attributes you perceive for the mead.

---

#### SOUR MEAD 1 Yeast: YH25

##### BOUQUET/AROMA

| Descriptors | Esters | Other | Comments |
|---|---|---|---|
| ☐ Delicate | ☐ Fruity | ☐ Bitter | |
| ☐ Light | ☐ Banana | ☐ Buttery | |
| ☐ Mild | ☐ Berry | ☐ Earthy | |
| ☑ Pleasant | ☐ Citrus | ☐ Grassy | |
| ☐ Rich | ☑ Tropical | ☑ Oaky | |
| ☐ Strong | ☐ Stone Fruit | ☑ Smoky | |
| ☐ Pungent | ☐ Unripe Fruit | ☑ Vegetal | |
| ☐ Sharp | ☐ Ripe Fruit | ☐ Vinous | |
| ☐ ________ | ☐ ________ | ☐ ________ | |

##### FLAVORS

| Descriptors | Esters | Other | Comments |
|---|---|---|---|
| ☐ Delicate | ☐ Apple | ☐ Bitter | |
| ☐ Light | ☐ Banana | ☑ Buttery | |
| ☐ Mild | ☐ Berry | ☐ Earthy | |
| ☑ Pleasant | ☐ Citrus | ☐ Grassy | |
| ☑ Rich | ☐ Fruity | ☐ Oaky | |
| ☐ Strong | ☐ Pear | ☐ Smoky | |
| ☐ Pungent | ☐ Stone Fruit | ☐ Sour | |
| ☐ Sharp | ☐ Tropical | ☐ Vegetal | |
| ☐ ________ | ☐ ________ | ☐ Vinous | |
| ☐ ________ | ☐ ________ | ☐ ________ | |

If this yeast were commercially available, would you buy it? ☐ No ☑ Maybe ☐ Absolutely

**OVERALL IMPRESSION**

☐ Yuck! ☐ Meh! ☑ Like it! ☐ Love it! ☐ Got to have it!

#### AMMA Sour Mead Tasting Feed Back Sheet

The purpose of this session is to gather as much feedback as possible on the **FLAVOR** profiles and attributes for the 13 different bacteria free, souring yeast strains used in these meads.

As a taster, do you consider yourself ☐ Novice ☐ Experienced ☐ Seasoned ☑ Professional

Check as many of the boxes for any **Bouquet/Aroma** (Smell) and **Flavor** (Taste) attributes you perceive for the mead.

---

#### SOUR MEAD 1 Yeast: YH25

**BOUQUET/AROMA**

| Descriptors | Esters | Other | Comments |
|---|---|---|---|
| ☑ Delicate | ☐ Fruity | ☐ Bitter | |
| ☑ Light | ☐ Banana | ☐ Buttery | |
| ☐ Mild | ☐ Berry | ☐ Earthy | |
| ☐ Pleasant | ☐ Citrus | ☐ Grassy | |
| ☐ Rich | ☐ Tropical | ☐ Oaky | |
| ☐ Strong | ☐ Stone Fruit | ☐ Smoky | |
| ☐ Pungent | ☐ Unripe Fruit | ☐ Vegetal | |
| ☐ Sharp | ☐ Ripe Fruit | ☐ Vinous ? | |
| ☐ ______ | ☑ Clean | ☐ ______ | |

**FLAVORS**

| Descriptors | Esters | Other | Comments |
|---|---|---|---|
| ☐ Delicate | ☐ Apple | ☐ Bitter | gross |
| ☐ Light | ☐ Banana | ☐ Buttery | |
| ☐ Mild | ☐ Berry | ☑ Earthy | |
| ☐ Pleasant | ☐ Citrus | ☐ Grassy | |
| ☐ Rich | ☐ Fruity | ☐ Oaky | |
| ☐ Strong | ☐ Pear | ☐ Smoky | |
| ☑ Pungent | ☐ Stone Fruit | ☐ Sour | |
| ☐ Sharp | ☐ Tropical | ☐ Vegetal | |
| ☐ ______ | ☐ Bretty | ☐ Vinous | |
| ☐ ______ | ☐ ______ | ☐ ______ | |

If this yeast were commercially available, would you buy it? ☐ No ☐ Maybe ☐ Absolutely

**OVERALL IMPRESSION**

☑ Yuck! ☐ Meh! ☐ Like it! ☐ Love it! ☐ Got to have it!

#### AMMA Sour Mead Tasting Feed Back Sheet

The purpose of this session is to gather as much feedback as possible on the **FLAVOR** profiles and attributes for the 13 different bacteria free, souring yeast strains used in these meads.

As a taster, do you consider yourself ☐ Novice ☐ Experienced ☒ Seasoned ☐ Professional

Check as many of the boxes for any **Bouquet/Aroma** (Smell) and **Flavor** (Taste) attributes you perceive for the mead.

---

#### SOUR MEAD 1 Yeast: YH25

**BOUQUET/AROMA**

| Descriptors | Esters | Other | Comments |
|---|---|---|---|
| ☐ Delicate | ☒ Fruity | ☐ Bitter | |
| ☐ Light | ☐ Banana | ☐ Buttery | |
| ☒ Mild | ☐ Berry | ☐ Earthy | |
| ☒ Pleasant | ☒ Citrus | ☐ Grassy | |
| ☐ Rich | ☒ Tropical | ☐ Oaky | |
| ☐ Strong | ☐ Stone Fruit | ☐ Smoky | |
| ☐ Pungent | ☐ Unripe Fruit | ☐ Vegetal | |
| ☐ Sharp | ☒ Ripe Fruit | ☐ Vinous | |
| ☐ ________ | ☐ ________ | ☐ ________ | |

**FLAVORS**

| Descriptors | Esters | Other | Comments |
|---|---|---|---|
| ☐ Delicate | ☐ Apple | ☐ Bitter | |
| ☐ Light | ☐ Banana | ☐ Buttery | |
| ☐ Mild | ☐ Berry | ☒ Earthy | |
| ☐ Pleasant | ☐ Citrus | ☐ Grassy | |
| ☒ Rich | ☒ Fruity | ☐ Oaky | |
| ☐ Strong | ☒ Pear | ☐ Smoky | |
| ☐ Pungent | ☐ Stone Fruit | ☐ Sour | |
| ☐ Sharp | ☒ Tropical | ☐ Vegetal | |
| ☐ ________ | ☐ ________ | ☐ Vinous | |
| ☐ ________ | ☐ ________ | ☒ Sweet | |

If this yeast were commercially available, would you buy it? ☒ No ☐ Maybe ☐ Absolutely

**OVERALL IMPRESSION**

☐ Yuck! ☒ Meh! ☐ Like it! ☐ Love it! ☐ Got to have it!

#### AMMA Sour Mead Tasting Feed Back Sheet

The purpose of this session is to gather as much feedback as possible on the **FLAVOR** profiles and attributes for the 13 different bacteria free, souring yeast strains used in these meads.

As a taster, do you consider yourself ☐ Novice ☐ Experienced ☒ Seasoned ☐ Professional

Check as many of the boxes for any **Bouquet/Aroma** (Smell) and **Flavor** (Taste) attributes you perceive for the mead.

---

#### SOUR MEAD 1 Yeast: YH25

##### BOUQUET/AROMA

| Descriptors | Esters | Other | Comments |
|---|---|---|---|
| ☐ Delicate | ☐ Fruity | ☐ Bitter | |
| ☒ Light | ☐ Banana | ☐ Buttery | |
| ☐ Mild | ☐ Berry | ☐ Earthy | |
| ☒ Pleasant | ☒ Citrus | ☐ Grassy | |
| ☐ Rich | ☒ Tropical | ☐ Oaky | |
| ☐ Strong | ☐ Stone Fruit | ☐ Smoky | |
| ☐ Pungent | ☐ Unripe Fruit | ☐ Vegetal | |
| ☐ Sharp | ☒ Ripe Fruit | ☐ Vinous | |
| ☐ ________ | ☐ ________ | ☐ ________ | |

##### FLAVORS

| Descriptors | Esters | Other | Comments |
|---|---|---|---|
| ☐ Delicate | ☐ Apple | ☐ Bitter | tastes like a Moscato |
| ☐ Light | ☐ Banana | ☐ Buttery | |
| ☐ Mild | ☐ Berry | ☐ Earthy | |
| ☒ Pleasant | ☐ Citrus | ☐ Grassy | |
| ☐ Rich | ☐ Fruity | ☐ Oaky | |
| ☐ Strong | ☒ Pear | ☐ Smoky | |
| ☐ Pungent | ☐ Stone Fruit | ☐ Sour | |
| ☐ Sharp | ☒ Tropical | ☐ Vegetal | |
| ☐ ________ | ☐ ________ | ☐ Vinous | |
| ☐ ________ | ☐ ________ | ☐ ________ | |

If this yeast were commercially available, would you buy it? ☐ No ☐ Maybe ☒ Absolutely

##### OVERALL IMPRESSION

☐ Yuck! ☐ Meh! ☐ Like it! ☐ Love it! ☒ Got to have it!

### AMMA Sour Mead Tasting Feed Back Sheet

The purpose of this session is to gather as much feedback as possible on the **FLAVOR** profiles and attributes for the 13 different bacteria free, souring yeast strains used in these meads.

As a taster, do you consider yourself ☐ Novice ☒ Experienced ☐ Seasoned ☐ Professional

Check as many of the boxes for any **Bouquet/Aroma** (Smell) and **Flavor** (Taste) attributes you perceive for the mead.

---

#### SOUR MEAD 1 Yeast: YH25

##### BOUQUET/AROMA

| Descriptors | Esters | Other |
|---|---|---|
| ☐ Delicate | ☒ Fruity | ☐ Bitter |
| ☒ Light | ☐ Banana | ☐ Buttery |
| ☐ Mild | ☐ Berry | ☐ Earthy |
| ☐ Pleasant | ☒ Citrus | ☐ Grassy |
| ☐ Rich | ☒ Tropical | ☐ Oaky |
| ☐ Strong | ☐ Stone Fruit | ☐ Smoky |
| ☐ Pungent | ☐ Unripe Fruit | ☐ Vegetal |
| ☐ Sharp | ☐ Ripe Fruit | ☐ Vinous |
| ☐ ________ | ☐ ________ | ☐ ________ |

**Comments**

Very light fruit, Citrus

##### FLAVORS

| Descriptors | Esters | Other |
|---|---|---|
| ☐ Delicate | ☐ Apple | ☐ Bitter |
| ☐ Light | ☐ Banana | ☐ Buttery |
| ☐ Mild | ☐ Berry | ☒ Earthy |
| ☒ Pleasant | ☒ Citrus | ☐ Grassy |
| ☐ Rich | ☒ Fruity | ☐ Oaky |
| ☐ Strong | ☐ Pear | ☐ Smoky |
| ☐ Pungent | ☐ Stone Fruit | ☒ Sour |
| ☐ Sharp | ☐ Tropical | ☐ Vegetal |
| ☐ ________ | ☐ ________ | ☐ Vinous |
| ☐ ________ | ☐ ________ | ☒ leather |

**Comments**

Tropical Fruit notes apparent with a leather Finish

If this yeast were commercially available, would you buy it? ☐ No ☒ Maybe ☐ Absolutely

##### OVERALL IMPRESSION

☐ Yuck! ☐ Meh! ☒ Like it! ☐ Love it! ☐ Got to have it!

#### AMMA Sour Mead Tasting Feed Back Sheet

The purpose of this session is to gather as much feedback as possible on the **FLAVOR** profiles and attributes for the 13 different bacteria free, souring yeast strains used in these meads.

As a taster, do you consider yourself ☐ Novice ☐ Experienced ☐ Seasoned ☒ Professional

Check as many of the boxes for any **Bouquet/Aroma** (Smell) and **Flavor** (Taste) attributes you perceive for the mead.

---

#### SOUR MEAD 1 Yeast: YH25

##### BOUQUET/AROMA

| Descriptors | Esters | Other | Comments |
|---|---|---|---|
| ☐ Delicate | ☐ Fruity | ☐ Bitter | |
| ☒ Light | ☐ Banana | ☒ Buttery | |
| ☐ Mild | ☐ Berry | ☐ Earthy | |
| ☐ Pleasant | ☒ Citrus | ☐ Grassy | |
| ☐ Rich | ☐ Tropical | ☐ Oaky | |
| ☐ Strong | ☐ Stone Fruit | ☐ Smoky | |
| ☐ Pungent | ☐ Unripe Fruit | ☐ Vegetal | |
| ☐ Sharp | ☐ Ripe Fruit | ☐ Vinous | |
| ☐ ________ | ☐ ________ | ☐ ________ | |

##### FLAVORS

| Descriptors | Esters | Other | Comments |
|---|---|---|---|
| ☐ Delicate | ☐ Apple | ☐ Bitter | |
| ☐ Light | ☐ Banana | ☒ Buttery | |
| ☐ Mild | ☐ Berry | ☐ Earthy | |
| ☒ Pleasant | ☒ Citrus | ☐ Grassy | |
| ☐ Rich | ☐ Fruity | ☐ Oaky | |
| ☐ Strong | ☐ Pear | ☐ Smoky | |
| ☐ Pungent | ☐ Stone Fruit | ☐ Sour | |
| ☐ Sharp | ☐ Tropical | ☐ Vegetal | |
| ☐ ________ | ☐ ________ | ☐ Vinous | |
| ☐ ________ | ☐ ________ | ☐ ________ | |

If this yeast were commercially available, would you buy it? ☐ No ☐ Maybe ☐ Absolutely

##### OVERALL IMPRESSION

☐ Yuck! ☐ Meh! ☒ Like it! ☐ Love it! ☐ Got to have it!

#### AMMA Sour Mead Tasting Feed Back Sheet

The purpose of this session is to gather as much feedback as possible on the **FLAVOR** profiles and attributes for the 13 different bacteria free, souring yeast strains used in these meads.

As a taster, do you consider yourself ☐ Novice ☒ Experienced ☐ Seasoned ☐ Professional

Check as many of the boxes for any **Bouquet/Aroma** (Smell) and **Flavor** (Taste) attributes you perceive for the mead.

---

#### SOUR MEAD 1 Yeast: YH25

##### BOUQUET/AROMA

| Descriptors | Esters | Other | Comments |
|---|---|---|---|
| ☒ Delicate | ☐ Fruity | ☐ Bitter | |
| ☐ Light | ☐ Banana | ☐ Buttery | |
| ☐ Mild | ☐ Berry | ☐ Earthy | |
| ☐ Pleasant | ☐ Citrus | ☐ Grassy | |
| ☐ Rich | ☐ Tropical | ☐ Oaky | |
| ☐ Strong | ☐ Stone Fruit | ☐ Smoky | |
| ☐ Pungent | ☐ Unripe Fruit | ☐ Vegetal | |
| ☐ Sharp | ☐ Ripe Fruit | ☐ Vinous | |
| ☐ ________ | ☐ ________ | ☐ ________ | |

##### FLAVORS

| Descriptors | Esters | Other | Comments |
|---|---|---|---|
| ☒ Delicate | ☐ Apple | ☐ Bitter | |
| ☐ Light | ☐ Banana | ☐ Buttery | |
| ☐ Mild | ☐ Berry | ☒ Earthy | |
| ☐ Pleasant | ☐ Citrus | ☒ Grassy | |
| ☐ Rich | ☐ Fruity | ☐ Oaky | |
| ☐ Strong | ☐ Pear | ☐ Smoky | |
| ☐ Pungent | ☐ Stone Fruit | ☐ Sour | |
| ☐ Sharp | ☐ Tropical | ☒ Vegetal | |
| ☐ ________ | ☐ ________ | ☐ Vinous | |
| ☐ ________ | ☐ ________ | ☐ ________ | |

If this yeast were commercially available, would you buy it? ☒ No ☐ Maybe ☐ Absolutely

##### OVERALL IMPRESSION

☐ Yuck! ☒ Meh! ☐ Like it! ☐ Love it! ☐ Got to have it!

### AMMA Sour Mead Tasting Feed Back Sheet

The purpose of this session is to gather as much feedback as possible on the **FLAVOR** profiles and attributes for the 13 different bacteria free, souring yeast strains used in these meads.

As a taster, do you consider yourself  [ ] Novice  [x] Experienced  [ ] Seasoned  [ ] Professional

Check as many of the boxes for any **Bouquet/Aroma** (Smell) and **Flavor** (Taste) attributes you perceive for the mead.

---

## SOUR MEAD 1   Yeast: YH25

##### BOUQUET/AROMA

| Descriptors | Esters | Other | Comments |
|---|---|---|---|
| [ ] Delicate | [x] Fruity | [ ] Bitter | |
| [ ] Light | [ ] Banana | [ ] Buttery | |
| [ ] Mild | [ ] Berry | [ ] Earthy | |
| [ ] Pleasant | [ ] Citrus | [ ] Grassy | |
| [ ] Rich | [x] Tropical | [ ] Oaky | |
| [ ] Strong | [x] Stone Fruit | [ ] Smoky | |
| [ ] Pungent | [ ] Unripe Fruit | [ ] Vegetal | |
| [x] Sharp | [ ] Ripe Fruit | [ ] Vinous | |
| [ ] ________ | [ ] ________ | [ ] ________ | |

##### FLAVORS

| Descriptors | Esters | Other | Comments |
|---|---|---|---|
| [ ] Delicate | [ ] Apple | [ ] Bitter | Pineapple |
| [ ] Light | [ ] Banana | [ ] Buttery | |
| [x] Mild | [ ] Berry | [ ] Earthy | |
| [ ] Pleasant | [x] Citrus | [ ] Grassy | |
| [ ] Rich | [x] Fruity | [ ] Oaky | |
| [ ] Strong | [ ] Pear | [ ] Smoky | |
| [ ] Pungent | [ ] Stone Fruit | [ ] Sour | |
| [ ] Sharp | [x] Tropical | [ ] Vegetal | |
| [ ] ________ | [ ] ________ | [ ] Vinous | |
| [ ] ________ | [ ] ________ | [ ] ________ | |

If this yeast were commercially available, would you buy it?  [ ] No  [x] Maybe  [ ] Absolutely

##### OVERALL IMPRESSION

[ ] Yuck!  [ ] Meh!  [x] Like it!  [ ] Love it!  [ ] Got to have it!

### AMMA Sour Mead Tasting Feed Back Sheet

The purpose of this session is to gather as much feedback as possible on the **FLAVOR** profiles and attributes for the 13 different bacteria free, souring yeast strains used in these meads.

As a taster, do you consider yourself [x] Novice ☐ Experienced ☐ Seasoned ☐ Professional

Check as many of the boxes for any **Bouquet/Aroma** (Smell) and **Flavor** (Taste) attributes you perceive for the mead.

---

#### SOUR MEAD 1 Yeast: YH25

##### BOUQUET/AROMA

| Descriptors | Esters | Other | Comments |
|---|---|---|---|
| ☐ Delicate | [x] Fruity | ☐ Bitter | |
| ☐ Light | ☐ Banana | ☐ Buttery | |
| [x] Mild | ☐ Berry | ☐ Earthy | |
| ☐ Pleasant | ☐ Citrus | ☐ Grassy | |
| ☐ Rich | ☐ Tropical | ☐ Oaky | |
| ☐ Strong | ☐ Stone Fruit | ☐ Smoky | |
| ☐ Pungent | [x] Unripe Fruit | [x] Vegetal | |
| ☐ Sharp | ☐ Ripe Fruit | ☐ Vinous | |
| ☐ ________ | ☐ ________ | ☐ ________ | |

##### FLAVORS

| Descriptors | Esters | Other | Comments |
|---|---|---|---|
| ☐ Delicate | ☐ Apple | ☐ Bitter | |
| ☐ Light | [x] Banana | [x] Buttery | |
| ☐ Mild | ☐ Berry | ☐ Earthy | |
| ☐ Pleasant | ☐ Citrus | ☐ Grassy | |
| ☐ Rich | [x] Fruity | ☐ Oaky | |
| [x] Strong | ☐ Pear | ☐ Smoky | |
| ☐ Pungent | ☐ Stone Fruit | ☐ Sour | |
| ☐ Sharp | ☐ Tropical | ☐ Vegetal | |
| ☐ ________ | ☐ ________ | ☐ Vinous | |
| ☐ ________ | ☐ ________ | ☐ ________ | |

If this yeast were commercially available, would you buy it? [x] No ☐ Maybe ☐ Absolutely

##### OVERALL IMPRESSION

☐ Yuck! [x] Meh! ☐ Like it! ☐ Love it! ☐ Got to have it!

#### AMMA Sour Mead Tasting Feed Back Sheet

The purpose of this session is to gather as much feedback as possible on the **FLAVOR** profiles and attributes for the 13 different bacteria free, souring yeast strains used in these meads.

As a taster, do you consider yourself ☐ Novice ☒ Experienced ☐ Seasoned ☐ Professional

Check as many of the boxes for any **Bouquet/Aroma** (Smell) and **Flavor** (Taste) attributes you perceive for the mead.

---

#### SOUR MEAD 1 Yeast: YH25

##### BOUQUET/AROMA

| Descriptors | Esters | Other |
|---|---|---|
| ☒ Delicate | ☒ Fruity | ☐ Bitter |
| ☐ Light | ☐ Banana | ☐ Buttery |
| ☐ Mild | ☐ Berry | ☐ Earthy |
| ☒ Pleasant | ☐ Citrus | ☐ Grassy |
| ☐ Rich | ☐ Tropical | ☐ Oaky |
| ☐ Strong | ☐ Stone Fruit | ☐ Smoky |
| ☐ Pungent | ☐ Unripe Fruit | ☐ Vegetal |
| ☐ Sharp | ☐ Ripe Fruit | ☐ Vinous |
| ☐ ________ | ☐ ________ | ☐ ________ |

**Comments**

smell is fruity and pleasant

##### FLAVORS

| Descriptors | Esters | Other |
|---|---|---|
| ☒ Delicate | ☒ Apple | ☐ Bitter |
| ☒ Light | ☐ Banana | ☐ Buttery |
| ☐ Mild | ☐ Berry | ☐ Earthy |
| ☐ Pleasant | ☐ Citrus | ☐ Grassy |
| ☐ Rich | ☒ Fruity | ☐ Oaky |
| ☐ Strong | ☒ Pear | ☐ Smoky |
| ☐ Pungent | ☐ Stone Fruit | ☐ Sour |
| ☐ Sharp | ☐ Tropical | ☐ Vegetal |
| ☐ ________ | ☐ ________ | ☐ Vinous |
| ☐ ________ | ☐ ________ | ☐ ________ |

**Comments**

not very sour at all
nice light, fruity flavors
maybe a mix of apple & pear

If this yeast were commercially available, would you buy it? ☐ No ☐ Maybe ☒ Absolutely

##### OVERALL IMPRESSION

☐ Yuck! ☐ Meh! ☐ Like it! ☒ Love it! ☐ Got to have it!

#### AMMA Sour Mead Tasting Feed Back Sheet

The purpose of this session is to gather as much feedback as possible on the **FLAVOR** profiles and attributes for the 13 different bacteria free, souring yeast strains used in these meads.

As a taster, do you consider yourself ☐ Novice ☐ Experienced ☐ Seasoned ☐ Professional

Check as many of the boxes for any **Bouquet/Aroma** (Smell) and **Flavor** (Taste) attributes you perceive for the mead.

---

#### SOUR MEAD 1 Yeast: YH25

##### BOUQUET/AROMA

| Descriptors | Esters | Other |
|---|---|---|
| ☐ Delicate | ☐ Fruity | ☐ Bitter |
| ☐ Light | ☐ Banana | ☒ Buttery |
| ☐ Mild | ☐ Berry | ☐ Earthy |
| ☐ Pleasant | ☐ Citrus | ☐ Grassy |
| ☐ Rich | ☐ Tropical | ☐ Oaky |
| ☐ Strong | ☐ Stone Fruit | ☐ Smoky |
| ☐ Pungent | ☐ Unripe Fruit | ☒ Vegetal |
| ☐ Sharp | ☐ Ripe Fruit | ☐ Vinous |
| ☐ ________ | ☐ ________ | ☐ ________ |

**Comments**

##### FLAVORS

| Descriptors | Esters | Other |
|---|---|---|
| ☐ Delicate | ☐ Apple | ☐ Bitter |
| ☐ Light | ☐ Banana | ☐ Buttery |
| ☐ Mild | ☐ Berry | ☐ Earthy |
| ☐ Pleasant | ☐ Citrus | ☐ Grassy |
| ☐ Rich | ☐ Fruity | ☐ Oaky |
| ☐ Strong | ☒ Pear | ☐ Smoky |
| ☐ Pungent | ☐ Stone Fruit | ☐ Sour |
| ☐ Sharp | ☐ Tropical | ☐ Vegetal |
| ☐ ________ | ☐ ________ | ☐ Vinous |
| ☐ ________ | ☐ ________ | ☐ ________ |

**Comments**

toast [illegible], not sour particularly
salty finish lingers

If this yeast were commercially available, would you buy it? ☐ No ☐ Maybe ☐ Absolutely

##### OVERALL IMPRESSION

☐ Yuck! ☒ Meh! ☐ Like it! ☐ Love it! ☐ Got to have it!

#### AMMA Sour Mead Tasting Feed Back Sheet

The purpose of this session is to gather as much feedback as possible on the **FLAVOR** profiles and attributes for the 13 different bacteria free, souring yeast strains used in these meads.

As a taster, do you consider yourself ☐ Novice ☐ Experienced ☐ Seasoned ☒ Professional

Check as many of the boxes for any **Bouquet/Aroma** (Smell) and **Flavor** (Taste) attributes you perceive for the mead.

---

#### SOUR MEAD 1 Yeast: YH25

##### BOUQUET/AROMA

| Descriptors | Esters | Other | Comments |
|---|---|---|---|
| ☒ Delicate | ☒ Fruity | ☐ Bitter | |
| ☒ Light | ☒ Banana | ☒ Buttery | |
| ☒ Mild | ☐ Berry | ☐ Earthy | |
| ☒ Pleasant | ☒ Citrus | ☐ Grassy | |
| ☐ Rich | ☐ Tropical | ☐ Oaky | |
| ☐ Strong | ☐ Stone Fruit | ☐ Smoky | |
| ☐ Pungent | ☐ Unripe Fruit | ☐ Vegetal | |
| ☐ Sharp | ☒ Ripe Fruit | ☐ Vinous | |
| ☐ ________ | ☐ ________ | ☐ ________ | |

##### FLAVORS

| Descriptors | Esters | Other | Comments |
|---|---|---|---|
| ☒ Delicate | ☒ Apple | ☐ Bitter | |
| ☒ Light | ☒ Banana | ☒ Buttery | |
| ☐ Mild | ☐ Berry | ☐ Earthy | |
| ☒ Pleasant | ☒ Citrus | ☐ Grassy | |
| ☐ Rich | ☒ Fruity | ☐ Oaky | |
| ☐ Strong | ☒ Pear | ☐ Smoky | |
| ☐ Pungent | ☐ Stone Fruit | ☒ Sour | |
| ☐ Sharp | ☒ Tropical | ☐ Vegetal | |
| ☐ ________ | ☐ ________ | ☐ Vinous | |
| ☐ ________ | ☐ ________ | ☐ ________ | |

If this yeast were commercially available, would you buy it? ☐ No ☐ Maybe ☐ Absolutely

##### OVERALL IMPRESSION

☐ Yuck! ☐ Meh! ☐ Like it! ☒ Love it! ☐ Got to have it!

*Could drink the whole glass. Tropical lemon drop. Totally want to pair this with tropical salsas on grilled chicken + fish.*

### AMMA Sour Mead Tasting Feed Back Sheet

The purpose of this session is to gather as much feedback as possible on the **FLAVOR** profiles and attributes for the 13 different bacteria free, souring yeast strains used in these meads.

As a taster, do you consider yourself ☑ Novice ☐ Experienced ☐ Seasoned ☐ Professional

Check as many of the boxes for any **Bouquet/Aroma** (Smell) and **Flavor** (Taste) attributes you perceive for the mead.

---

#### SOUR MEAD 1 Yeast: YH25

##### BOUQUET/AROMA

| Descriptors | Esters | Other | Comments |
|---|---|---|---|
| ☐ Delicate | ☐ Fruity | ☐ Bitter | |
| ☐ Light | ☐ Banana | ☐ Buttery | |
| ☐ Mild | ☐ Berry | ☐ Earthy | |
| ☑ Pleasant | ☐ Citrus | ☐ Grassy | |
| ☐ Rich | ☐ Tropical | ☐ Oaky | |
| ☐ Strong | ☐ Stone Fruit | ☐ Smoky | |
| ☐ Pungent | ☐ Unripe Fruit | ☐ Vegetal | |
| ☐ Sharp | ☐ Ripe Fruit | ☐ Vinous | |
| ☐ ________ | ☐ ________ | ☐ ________ | |

##### FLAVORS

| Descriptors | Esters | Other | Comments |
|---|---|---|---|
| ☐ Delicate | ☐ Apple | ☐ Bitter | |
| ☐ Light | ☐ Banana | ☐ Buttery | |
| ☐ Mild | ☐ Berry | ☐ Earthy | |
| ☑ Pleasant | ☐ Citrus | ☐ Grassy | |
| ☐ Rich | ☐ Fruity | ☐ Oaky | |
| ☐ Strong | ☑ Pear | ☐ Smoky | |
| ☐ Pungent | ☐ Stone Fruit | ☐ Sour | |
| ☐ Sharp | ☐ Tropical | ☐ Vegetal | |
| ☐ ________ | ☐ ________ | ☐ Vinous | |
| ☐ ________ | ☐ ________ | ☐ Sweet | |

If this yeast were commercially available, would you buy it? ☐ No ☑ Maybe ☐ Absolutely

**OVERALL IMPRESSION**

☐ Yuck! ☐ Meh! ☐ Like it! ☐ Love it! ☐ Got to have it!

### AMMA Sour Mead Tasting Feed Back Sheet

The purpose of this session is to gather as much feedback as possible on the **FLAVOR** profiles and attributes for the 13 different bacteria free, souring yeast strains used in these meads.

As a taster, do you consider yourself ☒ Novice ☐ Experienced ☐ Seasoned ☐ Professional

Check as many of the boxes for any **Bouquet/Aroma** (Smell) and **Flavor** (Taste) attributes you perceive for the mead.

---

#### SOUR MEAD 1 Yeast: YH25

##### BOUQUET/AROMA

| Descriptors | Esters | Other | Comments |
|---|---|---|---|
| ☐ Delicate | ☐ Fruity | ☐ Bitter | |
| ☐ Light | ☐ Banana | ☐ Buttery | |
| ☐ Mild | ☐ Berry | ☐ Earthy | |
| ☐ Pleasant | ☐ Citrus | ☐ Grassy | |
| ☒ Rich | ☒ Tropical | ☐ Oaky | |
| ☐ Strong | ☐ Stone Fruit | ☐ Smoky | |
| ☐ Pungent | ☐ Unripe Fruit | ☒ Vegetal | |
| ☐ Sharp | ☐ Ripe Fruit | ☐ Vinous | |
| ☐ ______ | ☐ ______ | ☐ ______ | |

##### FLAVORS

| Descriptors | Esters | Other | Comments |
|---|---|---|---|
| ☐ Delicate | ☐ Apple | ☐ Bitter | |
| ☐ Light | ☐ Banana | ☐ Buttery | |
| ☐ Mild | ☐ Berry | ☐ Earthy | |
| ☐ Pleasant | ☐ Citrus | ☐ Grassy | |
| ☐ Rich | ☒ Fruity | ☐ Oaky | |
| ☒ Strong | ☐ Pear | ☐ Smoky | |
| ☐ Pungent | ☐ Stone Fruit | ☐ Sour | |
| ☐ Sharp | ☐ Tropical | ☐ Vegetal | |
| ☐ ______ | ☐ ______ | ☐ Vinous | |
| ☐ ______ | ☐ ______ | ☒ Sweet | |

If this yeast were commercially available, would you buy it? ☐ No ☒ Maybe ☐ Absolutely

##### OVERALL IMPRESSION

☐ Yuck! ☐ Meh! ☒ Like it! ☐ Love it! ☐ Got to have it!

### AMMA Sour Mead Tasting Feed Back Sheet

The purpose of this session is to gather as much feedback as possible on the **FLAVOR** profiles and attributes for the 13 different bacteria free, souring yeast strains used in these meads.

As a taster, do you consider yourself ☐ Novice ☐ Experienced ☐ Seasoned ☒ Professional

Check as many of the boxes for any **Bouquet/Aroma** (Smell) and **Flavor** (Taste) attributes you perceive for the mead.

---

#### SOUR MEAD 1 Yeast: YH25

##### BOUQUET/AROMA

| Descriptors | Esters | Other | Comments |
|---|---|---|---|
| ☐ Delicate | ☒ Fruity | ☐ Bitter | fermenting grape must |
| ☐ Light | ☐ Banana | ☒ Buttery | |
| ☒ Mild | ☐ Berry | ☒ Earthy | |
| ☐ Pleasant | ☒ Citrus | ☒ Grassy | |
| ☐ Rich | ☒ Tropical | ☐ Oaky | |
| ☐ Strong | ☐ Stone Fruit | ☐ Smoky | |
| ☐ Pungent | ☐ Unripe Fruit | ☐ Vegetal | |
| ☐ Sharp | ☐ Ripe Fruit | ☐ Vinous | |
| ☐ ________ | ☐ ________ | ☐ ________ | |

##### FLAVORS

| Descriptors | Esters | Other | Comments |
|---|---|---|---|
| ☐ Delicate | ☒ Apple | ☐ Bitter | nutty finish |
| ☐ Light | ☐ Banana | ☒ Buttery | |
| ☒ Mild | ☐ Berry | ☐ Earthy | |
| ☐ Pleasant | ☐ Citrus | ☐ Grassy | |
| ☐ Rich | ☒ Fruity | ☒ Oaky | |
| ☐ Strong | ☐ Pear | ☐ Smoky | |
| ☐ Pungent | ☐ Stone Fruit | ☒ Sour | |
| ☐ Sharp | ☒ Tropical | ☒ Vegetal | |
| ☐ ________ | ☐ ________ | ☐ Vinous | |
| ☐ ________ | ☐ ________ | ☐ ________ | |

If this yeast were commercially available, would you buy it? ☒ No ☐ Maybe ☐ Absolutely

**OVERALL IMPRESSION**

☐ Yuck! ☒ Meh! ☐ Like it! ☐ Love it! ☐ Got to have it!

#### AMMA Sour Mead Tasting Feed Back Sheet

The purpose of this session is to gather as much feedback as possible on the **FLAVOR** profiles and attributes for the 13 different bacteria free, souring yeast strains used in these meads.

As a taster, do you consider yourself ☐ Novice ☐ Experienced ☐ Seasoned ☐ Professional

Check as many of the boxes for any **Bouquet/Aroma** (Smell) and **Flavor** (Taste) attributes you perceive for the mead.

---

#### SOUR MEAD 1 Yeast: YH25

##### BOUQUET/AROMA

| Descriptors | Esters | Other | Comments |
|---|---|---|---|
| ☑ Delicate | ☑ Fruity | ☐ Bitter | |
| ☐ Light | ☐ Banana | ☐ Buttery | |
| ☐ Mild | ☐ Berry | ☐ Earthy | |
| ☐ Pleasant | ☐ Citrus | ☐ Grassy | |
| ☐ Rich | ☐ Tropical | ☐ Oaky | |
| ☐ Strong | ☐ Stone Fruit | ☐ Smoky | |
| ☐ Pungent | ☐ Unripe Fruit | ☐ Vegetal | |
| ☐ Sharp | ☐ Ripe Fruit | ☐ Vinous | |
| ☐ ________ | ☐ ________ | ☐ ________ | |

##### FLAVORS

| Descriptors | Esters | Other | Comments |
|---|---|---|---|
| ☐ Delicate | ☐ Apple | ☐ Bitter | |
| ☐ Light | ☐ Banana | ☐ Buttery | |
| ☐ Mild | ☐ Berry | ☐ Earthy | |
| ☐ Pleasant | ☐ Citrus | ☐ Grassy | |
| ☐ Rich | ☐ Fruity | ☐ Oaky | |
| ☐ Strong | ☐ Pear | ☐ Smoky | |
| ☐ Pungent | ☐ Stone Fruit | ☐ Sour | |
| ☐ Sharp | ☐ Tropical | ☐ Vegetal | |
| ☐ ________ | ☐ ________ | ☐ Vinous | |
| ☐ ________ | ☐ ________ | ☐ ________ | |

If this yeast were commercially available, would you buy it? ☐ No ☑ Maybe ☐ Absolutely

**OVERALL IMPRESSION**

☐ Yuck! ☐ Meh! ☑ Like it! ☐ Love it! ☐ Got to have it!

#### AMMA Sour Mead Tasting Feed Back Sheet

The purpose of this session is to gather as much feedback as possible on the **FLAVOR** profiles and attributes for the 13 different bacteria free, souring yeast strains used in these meads.

As a taster, do you consider yourself ☒ Novice ☐ Experienced ☐ Seasoned ☐ Professional

Check as many of the boxes for any **Bouquet/Aroma** (Smell) and **Flavor** (Taste) attributes you perceive for the mead.

---

#### SOUR MEAD 1 Yeast: YH25

**BOUQUET/AROMA**

| Descriptors | Esters | Other | Comments |
|---|---|---|---|
| ☐ Delicate | ☐ Fruity | ☐ Bitter | |
| ☒ Light | ☒ Banana | ☒ Buttery | |
| ☐ Mild | ☐ Berry | ☐ Earthy | |
| ☐ Pleasant | ☐ Citrus | ☐ Grassy | |
| ☐ Rich | ☐ Tropical | ☐ Oaky | |
| ☐ Strong | ☐ Stone Fruit | ☐ Smoky | |
| ☐ Pungent | ☐ Unripe Fruit | ☐ Vegetal | |
| ☐ Sharp | ☐ Ripe Fruit | ☐ Vinous | |
| ☐ ________ | ☐ ________ | ☐ ________ | |

**FLAVORS**

| Descriptors | Esters | Other | Comments |
|---|---|---|---|
| ☐ Delicate | ☐ Apple | ☐ Bitter | |
| ☐ Light | ☒ Banana | ☐ Buttery | |
| ☐ Mild | ☐ Berry | ☐ Earthy | |
| ☐ Pleasant | ☐ Citrus | ☐ Grassy | |
| ☒ Rich | ☐ Fruity | ☐ Oaky | |
| ☐ Strong | ☐ Pear | ☐ Smoky | |
| ☐ Pungent | ☐ Stone Fruit | ☐ Sour | |
| ☐ Sharp | ☐ Tropical | ☐ Vegetal | |
| ☐ ________ | ☐ ________ | ☐ Vinous | |
| ☐ ________ | ☐ ________ | ☒ Bread | |

If this yeast were commercially available, would you buy it? ☒ No ☐ Maybe ☐ Absolutely

**OVERALL IMPRESSION**

☐ Yuck! ☒ Meh! ☐ Like it! ☐ Love it! ☐ Got to have it!

#### AMMA Sour Mead Tasting Feed Back Sheet

The purpose of this session is to gather as much feedback as possible on the **FLAVOR** profiles and attributes for the 13 different bacteria free, souring yeast strains used in these meads.

As a taster, do you consider yourself ☑ Novice ☐ Experienced ☐ Seasoned ☐ Professional

Check as many of the boxes for any **Bouquet/Aroma** (Smell) and **Flavor** (Taste) attributes you perceive for the mead.

---

#### SOUR MEAD 1 Yeast: YH25

**BOUQUET/AROMA**

| Descriptors | Esters | Other | Comments |
|---|---|---|---|
| ☐ Delicate | ☐ Fruity | ☐ Bitter | |
| ☐ Light | ☐ Banana | ☐ Buttery | |
| ☐ Mild | ☐ Berry | ☐ Earthy | |
| ☑ Pleasant | ☐ Citrus | ☐ Grassy | |
| ☐ Rich | ☑ Tropical | ☐ Oaky | |
| ☐ Strong | ☐ Stone Fruit | ☐ Smoky | |
| ☐ Pungent | ☐ Unripe Fruit | ☐ Vegetal | |
| ☐ Sharp | ☐ Ripe Fruit | ☐ Vinous | |
| ☐ ______ | ☐ ______ | ☐ ______ | |

**FLAVORS**

| Descriptors | Esters | Other | Comments |
|---|---|---|---|
| ☐ Delicate | ☐ Apple | ☐ Bitter | |
| ☑ Light | ☐ Banana | ☑ Buttery | |
| ☐ Mild | ☐ Berry | ☐ Earthy | |
| ☑ Pleasant | ☐ Citrus | ☐ Grassy | |
| ☐ Rich | ☑ Fruity | ☐ Oaky | |
| ☐ Strong | ☑ Pear | ☐ Smoky | |
| ☐ Pungent | ☐ Stone Fruit | ☐ Sour | |
| ☐ Sharp | ☐ Tropical | ☐ Vegetal | |
| ☐ ______ | ☐ ______ | ☐ Vinous | |
| ☐ ______ | ☐ ______ | ☐ ______ | |

If this yeast were commercially available, would you buy it? ☐ No ☐ Maybe ☐ Absolutely

**OVERALL IMPRESSION**

☐ Yuck! ☐ Meh! ☐ Like it! ☑ Love it! ☐ Got to have it!

#### AMMA Sour Mead Tasting Feed Back Sheet

The purpose of this session is to gather as much feedback as possible on the **FLAVOR** profiles and attributes for the 13 different bacteria free, souring yeast strains used in these meads.

As a taster, do you consider yourself ☐ Novice ☑ Experienced ☐ Seasoned ☐ Professional

Check as many of the boxes for any **Bouquet/Aroma** (Smell) and **Flavor** (Taste) attributes you perceive for the mead.

---

#### SOUR MEAD 1 Yeast: YH25

**BOUQUET/AROMA**

| Descriptors | Esters | Other | Comments |
|---|---|---|---|
| ☐ Delicate | ☐ Fruity | ☐ Bitter | |
| ☐ Light | ☑ Banana | ☐ Buttery | |
| ☑ Mild | ☐ Berry | ☐ Earthy | |
| ☐ Pleasant | ☐ Citrus | ☐ Grassy | |
| ☐ Rich | ☐ Tropical | ☐ Oaky | |
| ☐ Strong | ☐ Stone Fruit | ☐ Smoky | |
| ☐ Pungent | ☐ Unripe Fruit | ☐ Vegetal | |
| ☐ Sharp | ☐ Ripe Fruit | ☐ Vinous | |
| ☐ ______ | ☐ ______ | ☐ acidic | |

**FLAVORS**

| Descriptors | Esters | Other | Comments |
|---|---|---|---|
| ☐ Delicate | ☐ Apple | ☐ Bitter | |
| ☐ Light | ☐ Banana | ☐ Buttery | |
| ☑ Mild | ☐ Berry | ☐ Earthy | |
| ☐ Pleasant | ☑ Citrus | ☐ Grassy | |
| ☐ Rich | ☐ Fruity | ☐ Oaky | |
| ☐ Strong | ☑ Pear | ☐ Smoky | |
| ☐ Pungent | ☐ Stone Fruit | ☐ Sour | |
| ☐ Sharp | ☐ Tropical | ☐ Vegetal | |
| ☐ ______ | ☐ ______ | ☐ Vinous | |
| ☐ ______ | ☐ ______ | ☐ ______ | |

If this yeast were commercially available, would you buy it? ☑ No ☐ Maybe ☐ Absolutely

**OVERALL IMPRESSION**

☐ Yuck! ☑ Meh! ☐ Like it! ☐ Love it! ☐ Got to have it!

#### AMMA Sour Mead Tasting Feed Back Sheet

The purpose of this session is to gather as much feedback as possible on the **FLAVOR** profiles and attributes for the 13 different bacteria free, souring yeast strains used in these meads.

As a taster, do you consider yourself ☐ Novice ☑ Experienced ☐ Seasoned ☐ Professional

Check as many of the boxes for any **Bouquet/Aroma** (Smell) and **Flavor** (Taste) attributes you perceive for the mead.

---

#### SOUR MEAD 1 Yeast: YH25

##### BOUQUET/AROMA

| Descriptors | Esters | Other | Comments |
|---|---|---|---|
| ☐ Delicate | ☐ Fruity | ☐ Bitter | ______ |
| ☑ Light | ☑ Banana | ☐ Buttery | ______ |
| ☐ Mild | ☐ Berry | ☐ Earthy | ______ |
| ☐ Pleasant | ☐ Citrus | ☐ Grassy | ______ |
| ☐ Rich | ☐ Tropical | ☑ Oaky | ______ |
| ☐ Strong | ☐ Stone Fruit | ☐ Smoky | ______ |
| ☐ Pungent | ☐ Unripe Fruit | ☐ Vegetal | ______ |
| ☐ Sharp | ☐ Ripe Fruit | ☐ Vinous | |
| ☐ ______ | ☐ ______ | ☐ ______ | |

##### FLAVORS

| Descriptors | Esters | Other | Comments |
|---|---|---|---|
| ☑ Delicate | ☐ Apple | ☐ Bitter | ______ |
| ☐ Light | ☐ Banana | ☐ Buttery | ______ |
| ☐ Mild | ☐ Berry | ☐ Earthy | ______ |
| ☐ Pleasant | ☐ Citrus | ☐ Grassy | ______ |
| ☐ Rich | ☑ Fruity | ☐ Oaky | ______ |
| ☐ Strong | ☐ Pear | ☐ Smoky | ______ |
| ☐ Pungent | ☐ Stone Fruit | ☐ Sour | ______ |
| ☐ Sharp | ☐ Tropical | ☐ Vegetal | ______ |
| ☐ ______ | ☐ ______ | ☐ Vinous | |
| ☐ ______ | ☐ ______ | ☐ ______ | |

If this yeast were commercially available, would you buy it? ☐ No ☑ Maybe ☐ Absolutely

##### OVERALL IMPRESSION

☐ Yuck! ☐ Meh! ☑ Like it! ☐ Love it! ☐ Got to have it!

#### AMMA Sour Mead Tasting Feed Back Sheet

The purpose of this session is to gather as much feedback as possible on the **FLAVOR** profiles and attributes for the 13 different bacteria free, souring yeast strains used in these meads.

As a taster, do you consider yourself ☐ Novice ☐ Experienced ☐ Seasoned ☒ Professional

Check as many of the boxes for any **Bouquet/Aroma** (Smell) and **Flavor** (Taste) attributes you perceive for the mead.

---

#### SOUR MEAD 1 Yeast: YH25

##### BOUQUET/AROMA

| Descriptors | Esters | Other | Comments |
|---|---|---|---|
| ☐ Delicate | ☒ Fruity | ☐ Bitter | |
| ☐ Light | ☐ Banana | ☐ Buttery | |
| ☒ Mild | ☐ Berry | ☐ Earthy | |
| ☐ Pleasant | ☐ Citrus | ☐ Grassy | |
| ☐ Rich | ☒ Tropical | ☐ Oaky | |
| ☐ Strong | ☐ Stone Fruit | ☒ Smoky | |
| ☐ Pungent | ☐ Unripe Fruit | ☐ Vegetal | |
| ☐ Sharp | ☐ Ripe Fruit | ☐ Vinous | |
| ☐ ________ | ☐ ________ | ☐ ________ | |

##### FLAVORS

| Descriptors | Esters | Other | Comments |
|---|---|---|---|
| ☐ Delicate | ☐ Apple | ☐ Bitter | |
| ☐ Light | ☐ Banana | ☐ Buttery | |
| ☐ Mild | ☐ Berry | ☐ Earthy | |
| ☐ Pleasant | ☐ Citrus | ☐ Grassy | |
| ☒ Rich | ☐ Fruity | ☐ Oaky | |
| ☐ Strong | ☒ Pear | ☒ Smoky | |
| ☐ Pungent | ☐ Stone Fruit | ☐ Sour | |
| ☐ Sharp | ☐ Tropical | ☐ Vegetal | |
| ☐ ________ | ☐ ________ | ☐ Vinous | |
| ☐ ________ | ☐ ________ | ☐ ________ | |

If this yeast were commercially available, would you buy it? ☐ No ☒ Maybe ☐ Absolutely

**OVERALL IMPRESSION**

☐ Yuck! ☐ Meh! ☒ Like it! ☐ Love it! ☐ Got to have it!

#### AMMA Sour Mead Tasting Feed Back Sheet

The purpose of this session is to gather as much feedback as possible on the **FLAVOR** profiles and attributes for the 13 different bacteria free, souring yeast strains used in these meads.

As a taster, do you consider yourself ☐ Novice ☐ Experienced ☒ Seasoned ☐ Professional

Check as many of the boxes for any **Bouquet/Aroma** (Smell) and **Flavor** (Taste) attributes you perceive for the mead.

---

#### SOUR MEAD 1 Yeast: YH25

##### BOUQUET/AROMA

| Descriptors | Esters | Other | Comments |
|---|---|---|---|
| ☒ Delicate | ☐ Fruity | ☐ Bitter | |
| ☐ Light | ☐ Banana | ☐ Buttery | |
| ☐ Mild | ☐ Berry | ☐ Earthy | |
| ☐ Pleasant | ☒ Citrus | ☐ Grassy | |
| ☐ Rich | ☐ Tropical | ☐ Oaky | |
| ☐ Strong | ☐ Stone Fruit | ☐ Smoky | |
| ☐ Pungent | ☐ Unripe Fruit | ☐ Vegetal | |
| ☐ Sharp | ☐ Ripe Fruit | ☐ Vinous | |
| ☐ ________ | ☐ ________ | ☐ ________ | |

##### FLAVORS

| Descriptors | Esters | Other | Comments |
|---|---|---|---|
| ☐ Delicate | ☒ Apple | ☐ Bitter | |
| ☐ Light | ☐ Banana | ☐ Buttery | |
| ☐ Mild | ☐ Berry | ☐ Earthy | |
| ☐ Pleasant | ☐ Citrus | ☐ Grassy | |
| ☐ Rich | ☐ Fruity | ☐ Oaky | |
| ☐ Strong | ☐ Pear | ☐ Smoky | |
| ☐ Pungent | ☐ Stone Fruit | ☒ Sour | |
| ☐ Sharp | ☐ Tropical | ☐ Vegetal | |
| ☐ ________ | ☐ ________ | ☐ Vinous | |
| ☐ ________ | ☐ ________ | ☐ ________ | |

If this yeast were commercially available, would you buy it? ☐ No ☐ Maybe ☐ Absolutely

**OVERALL IMPRESSION**

☐ Yuck! ☒ Meh! ☐ Like it! ☐ Love it! ☐ Got to have it!
